# ASD mutations in the ciliary gene *CEP41* impact development of projection neurons and interneurons in a human cortical organoid model

**DOI:** 10.1101/2025.07.09.663904

**Authors:** Kerstin Hasenpusch-Theil, Alexandra Lesayova, Zrinko Kozic, Mariana Beltran, Grace Wilson, Neil C Henderson, Owen Dando, Thomas Theil

## Abstract

Primary cilia control cell-cell signalling and their dysfunction has been implicated in Autism Spectrum Disorders (ASD) but their roles in the ASD aetiology remain largely unexplored. Here, we analysed the impact of ASD mutations in *CEP41* using human corticogenesis. *CEP41* encodes a centrosomal protein located at the basal body and the ciliary axoneme and is mutated in ASD individuals and in Joubert syndrome, a ciliopathy with high incidence of ASD. To gain insights into *CEP41*’s role in ASD aetiology, we characterised human cortical organoids carrying the *CEP41* R242H point mutations found in ASD individuals. This mutation did not interfere with CEP41’s ciliary localisation but cilia were shorter and had lower levels of tubulin polyglutamylation, which is indicative of altered cilia stability and signalling. Moreover, scRNAseq analyses revealed that the expression of several transcription factors with critical roles in interneuron development was altered in mutant interneurons and their progenitors. The *CEP41* mutation also caused decreased cortical progenitor proliferation and an augmented formation of upper layer cortical neurons. Taken together, these findings indicate that *CEP41* controls excitatory and inhibitory neuron differentiation, alterations in which might lead to an excitation/inhibition imbalance that is widely recognized as a convergent mechanism underlying neurodevelopmental disorders.

## INTRODUCTION

Autism spectrum disorders (ASD) comprises a complex neurodevelopmental condition that is characterized by difficulties in social cognition and communication, repetitive behaviours and hypersensitivity to external stimuli^1^. These core symptoms are associated with several comorbidities that may contribute to the high variability of ASD symptoms^2–4^. Post-mortem studies^5^ support a long-standing hypothesis that links ASD and other neurodevelopmental disorders (NDDs) with an imbalance between excitation and inhibition (E/I imbalance)^6,7^, whereby interneurons regulate cortical circuitry through their inhibitory effects. While research into NDDs has mostly focused on altered neuronal connectivity and circuitry, there is increasing evidence that NDD symptoms can originate from brain malformations occurring in the second trimester of foetal development^8,9^. In this period, neural stem cells generate excitatory glutamatergic projection neurons and inhibitory GABAergic interneurons in the dorsal and ventral telencephalon, respectively. Interneurons, in particular, form an enormous variety of cell types that is critical for the proper functioning of cortical circuitry. The specification of this interneuron diversity is under the control of Sonic hedgehog (SHH) signalling and a cascade of transcription factors^10^. After their birth, projection neurons migrate radially out of the germinal zones to form the cortical layers in an inside-out manner, while interneurons undergo complex tangential migrations from the ventral telencephalon to their target area in the cortex where they mature and integrate into cortical circuits^11^.

Primary cilia are cellular antennas present on most cells including neural progenitors and many neurons and act as signalling hubs in development and tissue homeostasis. Defects in the function and/or structure of primary cilia underlie a group of syndromes commonly referred to as ciliopathies^12^ that are characterized by pleiotropic clinical features. Many ciliopathy patients display severe neurological symptoms, most commonly ID and ASD^13^. In turn, a cell-based high-throughput screen indicated that diverse neuropsychiatric risk genes converge on primary cilia^14^. This idea is further corroborated by recent findings that several monogenetic neurodevelopmental syndromes, including Fragile X and Rett syndromes, result in altered ciliary structure and signalling^15–19^, but the mechanisms how ciliary impairments contribute to NDD pathogenesis remain largely unexplored.

To date, the most compelling evidence for ciliary roles in ASD aetiology stems from the identification of autism specific mutations in several ciliary genes^20–26^. One of these genes, *CEP41*, encodes a centrosomal protein located at the basal body and in the ciliary axoneme. It is essential for the transport of Tubulin Tyrosine Ligase Like 6 (TTLL6) into the cilium and hence tubulin polyglutamylation^27^. Homozygous *CEP41* null mutations cause Joubert Syndrome (JS)^27^, a genetically and phenotypically heterogeneous syndrome^28^ associated with intellectual disabilities (ID) and ASD in 40% of JS patients^20^. In addition, heterozygous *CEP41* missense mutations have been identified in familial and sporadic forms of ASD^23,25^ but the effects of these mutations on mammalian brain development have not been explored. Here, we report our findings on human cortical organoids carrying the *CEP41* R242H ASD mutation^25^. This mutation did not interfere with CEP41’s ciliary localisation but cilia were shorter and had lower levels of tubulin polyglutamylation indicative of altered cilia stability and signalling. Moreover, scRNA-seq analyses revealed that the *CEP41* mutation caused decreased cortical progenitor proliferation and an augmented formation of upper layer cortical neurons. In interneurons and their progenitors, the altered expression of several key transcription factors in interneuron development coincided with changes in interneuron differentiation. Taken together, these results indicate that the combined effects of the ASD-linked *CEP41* mutation on the development of projection neurons and interneurons may contribute to ASD pathogenesis.

## MATERIAL AND METHODS

### Cell culture

Feeder-free iPSCs were continuously maintained in StemMACS^TM^ iPS-Brew XF medium containing supplements (Miltenyi Biotec, #130-104-368) and 1x Antibiotic-Antimycotic (GIBCO, #15240062) on CellAdhere^TM^ Laminin-521 (STEMCELL Technologies UK Ltd; #77003) coated 6-well plates. All cells were maintained at 37°C in a 5% CO_2_ atmosphere.

### Gene editing by CRISPR/Cas9 Homology-Directed Repair

The *CEP41 R242H* mutation was generated in the Nas2 iPSC line^29^. To ensure the absence of unknown *CEP41* mutations, the targeted exon and flanking sequences were sequenced prior to gene editing using oligonucleotides CEP_E9_F and CEP_E9_R (Supplementary Table 1). The gRNA was designed using an online CRISPR design tool (http://crispr.mit.edu) and was cloned into the pSpCas9(BB)-2A-Puro (PX459) plasmid (Addgene: #48139). To test gRNA efficiency a T7 endonuclease assay was performed. Nas2 iPSC cells were cultured up to 70-80% confluency and lifted with Accutase enzyme (Invitrogen, #00-4555-56). gRNA constructs together with pmaxGFP^TM^ plasmid were transfected using a P3 Primary Cell 4D-Nucleofector Kit (Lonza, V4XP-3012) according to the manufacturer’s instructions. Cells were harvested 48 hours post-transfection and genomic DNA was extracted using the QuickExtract^TM^ DNA Extraction solution (LGC Biosearch Technologies, QE09050)). Genomic targeting efficiency for the gRNA was determined through annealing and digestion with T7 Endonuclease I (NEB: #M0302) of a PCR product flanking the *CEP41* R242H target site. gRNA 5’-GACCTCCGGAAAGCATGAAG-3’ was determined as optimal for use in gene-editing. iPSCs at 70-80% confluence were dissociated into single cells with Accutase and 8×10^5^ cells were electroporated with 5 μg Cas9-CEP41_sgRNA plasmid, 1.8 µg pmaxGFP^TM^ vector (Lonza kit) and 100 pmol 178nt Ultramer, a single-stranded DNA oligonucleotide donor template (ssODN) (PAGE-purified; Integrated DNA Technologies) (Supplementary Table 1), using the P3 Primary Cell 4D-Nucleofector^TM^ X Kit (Lonza), (program CA-137) on a Lonza 4D-Nucleofector™ X Unit (Lonza) according to manufacturer’s guidelines. Transfected cells were resuspended in pre-warmed StemMACS^TM^ iPS-Brew XF including supplements and 10 µM ROCK-inhibitor (Y-27632, Stemcell Technologies) and seeded into two wells of a Laminin-521 coated 6-well plate. Selection with 1 μg/ml puromycin (InvivoGen, #ant-pr-1) was commenced 24 hours post-nucleofection and continued for 24 hours. Cells were grown to confluence and passaged at low density (5×10^3^), as single cells onto Laminin-521 coated 10 cm dishes in StemMACS^TM^ iPS-Brew XF medium with 10 μM Y27632. After 4-6 days single-cell derived colonies were isolated and transferred to a Laminin-521 coated 96-well plate. Duplicate plates were made for maintenance and restriction fragment length polymorphism (RFLP) screening. When cells for genotyping reached confluence, crude genomic DNA lysates were prepared by adding 25μl QuickExtract^TM^ DNA Extraction solution and incubated at 65°C for 15 minutes, followed by 2 minutes at 98°C. Amplicons flanking the targeting site were amplified using GoTaq G2 polymerase (Promega) with the CEP_E9_F and CEP_E9_R primers (Supplementary Table 1). PCR protocol: 95_°_C for 2 minutes; 35 cycles of 95_°_C for 40 seconds, 57_°_C for 40 seconds, 72_°_C for 50 seconds; and a final extension at 72_°_C for 5 minutes. PCR products were digested with Bsp 1286I (New England Biolabs) and run on a 2.5% TAE agarose gel. Clones identified as carrying a Bsp 1286I restriction site were evaluated for introduction of the R242H mutation through Sanger sequencing (Source Bioscience). The top 5 candidates for off-target effects identified with an online tool (http://crispr.mit.edu) were sequenced using oligonucleotides as summarized in Supplementary Table 1. Successfully edited clones were expanded and assessed for karyotypes with the Aneuploidy kBoBs assay (TDL Genetics Ltd, London) (Supplementary Data). Quality control tests were performed after clonal passage 10 and included immunocytochemistry with a panel of antibodies to pluripotency markers.

### Generation of cerebral organoids

Cerebral organoids were generated and maintained according to a modified Lancaster protocol^30^ as described recently^31^ with media changes every second day. This protocol generates embryoid bodies for 6 days first before making neurospheres by dual-SMAD inhibition^32^. hiPSCs were cultured in CellAdhere^TM^ Laminin-521 coated 6-well plates in StemMACS^TM^ iPS-Brew XF including supplements and 1x Antibiotic-Antimycotic for an average of four to five days. When cultures reached around 80% confluency with distinct, well defined hiPSC colonies, cells were lifted with Accutase enzyme (Invitrogen, #00-4555-56) and resuspended in Stem StemMACS^TM^ iPS-Brew XF including supplements, 1x Antibiotic-Antimycotic, 50μM Rock Inhibitor Y-27632 dihydrochloride (TOCRIS, #1254) and 4ng/ml recombinant human FGF2 for four days (Peprotech, #100-18B). On day 6, embryoid bodies were transferred into Neural Induction media: 80% (v/v) DMEM/F-12, HEPES (GIBCO, #11330032), 20% (v/v) KnockOut^TM^ Serum Replacement (GIBCO, #10828010), 1x Antibiotic-Antimycotic (GIBCO, #15240062), 1x GlutaMAX^TM^ supplement (GIBCO, #35050061), 1x MEM-Non-Essential Amino Acids solution (GIBCO, #11140035), 0.1mM 2-Mercaptoethanol (GIBCO, #31350010), 10µM Activin Inhibitor SB 431542 (Tocris, #1614) and 0.1µM LDN 193189 (StemCell Technologies, #72147). From this point onwards, cells were cultured in suspension on an orbital shaker at 45 rpm in a cell culture CO_2_ incubator at 37⁰C and 5% CO_2_. After four days, colonies were transferred to EB1 medium containing Advanced DMEM/F-12 (GIBCO, #12634010) supplemented with 1x Antibiotic-Antimycotic, 1x GlutaMAX™Supplement, 1x N-2 Supplement (GIBCO, #17502048), 0.25x B-27 Supplement minus Vitamin A (GIBCO, #12587010) and 7μg/ml Heparin (StemCell Technologies, #07980). After 8 days, rosette forming spheres were transferred into EB2 medium until day 32. EB2 medium consisted of a 50% (v/v) Advanced DMEM/F-12, 50%(v/v) Neurobasal™ Medium (GIBCO, #21103049), supplemented with 1x Antibiotic-Antimycotic, 0.5x GlutaMAX™Supplement, 1x N-2 Supplement, 0.25x B-27 Supplement Minus Vitamin A, 1x MEM Non-Essential Amino Acids Solution, and 1.25 µg/ml human Insulin (Sigma-Aldrich, #I9278). At day 32, organoids were transferred into EB3 medium with 50% (v/v) Advanced DMEM/F-12, 50% (v/v) Neurobasal™ Medium, supplemented with 1x Antibiotic-Antimycotic, 1x GlutaMAX™Supplement, 0.5x N-2 Supplement, 0.5x B-27 Supplement (GIBCO, #17504044), 0.5x MEM Non-Essential Amino Acids Solution, 1mM 2-Mercaptoethanol, 2.5 µg/ml human Insulin for the remainder of organoid growth. 20ng/ml recombinant human/murine/rat BDNF (Peprotech, #450-02) and 20ng/ml recombinant human NT3 were added between days 32 and 48 (Peprotech, #450-03). At this time point, the speed of the orbital shaker was increased to 60 rpm. Organoids were collected at different developmental stages for immunohistochemistry, RNA or protein extraction.

### Generation of ventral telencephalic organoids

Ventral organoid differentiation was based on a protocol by the Pasca group^33^. iPSCs colonies were lifted with Accutase and resuspended in Stem StemMACS^TM^ iPS-Brew XF containing supplements, 1x Antibiotic-Antimycotic and 10μM Rock Inhibitor Y-27632 dihydrochloride (TOCRIS, #1254). 8000 cells were plated per well into a low adhesive 96 well-plate. On Day 1, medium was replenished before transferring spheroids on Day 2 into Neural induction medium: 80% (v/v) DMEM/F-12, HEPES, 20% (v/v) KnockOut^TM^ Serum Replacement, 1x Antibiotic-Antimycotic, 0.5x GlutaMAX^TM^ supplement, 1x MEM-Non-Essential Amino Acids solution, 0.1mM 2-Mercaptoethanol, 10µM Activin Inhibitor SB 431542 (Tocris, #1614) and 5µM Dorsomorphin (Sigma-Aldrich, #P5499) which was changed daily. For Day 5 and 6 the medium was supplemented with 5µM IWP-2 (LKT Laboratories, Inc., #I9060). After neural induction, organoids were transferred into Neuronal medium and placed into a 24 well-plate on an orbital shaker at 45rpm in a cell culture CO_2_ incubator at 37⁰C and 5% CO_2_ and fed every other day. Neuronal medium contained Neurobasal A medium (Life Technologies, 10888022), 1x Antibiotic-Antimycotic, 1xB-27 supplement without Vitamin A, 1x GlutaMAX™ Supplement. On Days 7–11, 20ng/ml recombinant human FGF-2 and 20ng/ml recombinant murine EGF (Peprotech, #315-09) were added. Day12–22 organoids were fed with neuronal medium containing 20ng/ml recombinant human FGF-2, 20ng/ml recombinant murine EGF and 100nM SAG (Cayman Chemical, #11914) to induce ventral telencephalic differentiation. On Day24, organoids received Neuronal medium only. From Day 25 until 43, Neuronal medium was supplemented with 20ng/ml recombinant human/murine/rat BDNF and 20ng/ml recombinant human NT3. Organoids were collected at different developmental stages for immunohistochemistry, RNA or protein extraction.

### Immunohistochemistry on organoids

For immunohistochemistry, organoids were fixed for 1 hour in 4% paraformaldehyde, incubated in 30% sucrose at +4°C for 24h, embedded in 30% sucrose/OCT mixture (1:1) and frozen on dry ice. Immunofluorescence staining was performed on 10-12 μm cryostat sections as described previously^34^ with antibodies against mouse anti-ARL13B (Neuromab 75-287; 1:2000), mouse anti-BrdU (Becton Dickinson #347580; 1:50), rabbit anti-CEP41 (Proteintech #17566-1-AP; 1:200), guinea pig anti-DLX2 (Bioacademica # 74-116; 1:2000), rabbit anti-pHH3 (Millipore #06-570; 1:100), rabbit anti-IFT88 (Proteintech #13967-1-AP; 1:200); rabbit anti-IFT144 (Proteintech #13647-1-AP; 1:200); mouse anti-NR2F2 (Persus Proteomics #PP-H7147-00; 1:300), rabbit anti-OLIG2 (Millipore #AB9610; 1:400), rabbit anti-PAX6 (Biolegend #901301; 1:400), mouse anti-SATB2 (Abcam #51502; 1:200), mouse anti-SOX2 (Santa Cruz Biotechnology, #sc-365823; 1:200), rabbit anti-SOX2 (Abcam #92494; 1:1000), rabbit anti-TBR1 (Abcam #31940; 1:400), mouse anti-γTUB (Sigma T6557; 1:2000), mouse anti-glutamylated TUBULIN GT335 (AdipoGen, #AG-20B-0020; 1:1000), mouse anti-pVIM (MBL, #D076-3; 1:500).

Primary antibodies for immunohistochemistry were detected with Alexa- or Cy2/3-conjugated fluorescent secondary antibodies. The TBR1 signal was amplified using biotinylated secondary IgG antibody (swine anti-rabbit IgG) (1:400, BD Biosciences) followed by Alexa Fluor 488 or 568 Streptavidin (1:100, Invitrogen). For counter staining DAPI (1:2000, Life Technologies) was used. Fluorescent and confocal images were taken on a LeicaDM 5500 B fluorescent microscope and Nikon A1R FLIM confocal microscope, respectively.

### Immunohistochemistry for pluripotency markers

hiPSCs were cultured in StemMACS^TM^ iPS-Brew XF containing supplements and 1x Antibiotic-Antimycotic on round glass coverslips in 24-well plates coated withCellAdhere^TM^ Laminin-521. Cells were grown for an average of 4-6 days until cultures reached around 80% confluency, when they were fixed for 15 min at room temperature in 4% paraformaldehyde/DPBS. To detect pluripotent specific antigens, the StemLight^TM^ Pluripotency Antibody Kit (Cell Signaling Technology, #9656) was used according to the manufacturer’s instructions with the following primary antibodies: rabbit anti-OCT4A, rabbit anti-SOX2, rabbit anti-NANOG, mouse anti-SSEA4, mouse anti-TRA-1-60(S) and mouse anti-TRA-1-81 (all 1:200). Primary antibodies were detected with Cy2-conjugated Donkey anti-rabbit IgG (1:100, Jackson ImmunoResearch, #711-225-152), Cy3-conjugated Donkey anti-mouse IgG (1:100; Jackson ImmunoResearch, #715-165-151) and Cy3-conjugated Donkey anti-mouse IgM (1:100, Jackson ImmunoResearch, #715-165-140) secondary antibodies. Cell nuclei were stained with DAPI ((1:2000, Invitrogen, #D1306). Fluorescent images were captures using a Leica DM5500 B fluorescent microscope.

### Western Blot

Protein was extracted from control, *CEP41*^R242H/+^ and *CEP41*^R242H/R242H^ organoids (derived from n=3 lines for each genotype) as described previously^35^. 30 μg protein lysates were subjected to gel electrophoresis on a 3-8% NuPAGE® Tris-Acetate gel (Life Technologies), and protein was transferred to an Immobilon-FL membrane (Millipore), which was incubated with rabbit anti-CEP41 (1:1000, Affinity Biosciences #DF9362) and mouse anti-β-GAPDH antibody (1:5000, Abcam #ab9484). After incubating with goat anti-rabbit IgG IRDye800CW (1:10,000, LI-COR Biosciences, #926-32211) and goat anti-mouse IgG Alexa Fluo 680 secondary antibodies (1:5000, Life Technologies, #A21058), signal was detected using the Odyssey M Imaging System (LICORbio) and LI-COR Acquisition 2.2 software. Values for protein signal intensity were obtained using Image Studio Lite Version 4.0 (LICORbio). CEP41 and GAPDH protein level ratios were compared between control and mutant organoids using an ordinary one-way ANOVA followed by Tukey’s multiple comparisons test.

### qRT-PCR

To validate differential expression of *GLI1*, total RNA was extracted from control and *CEP41*^R242H/R242H^ ventral telencephalic organoids (n=3 lines per genotype) using a RNeasy Plus Micro Kit (Qiagen) and reverse transcribed using Superscript™ IV VILO™ Master ezDNase enzyme (Thermo Fisher Scientific). Quantitative reverse transcription PCR (qRT-PCR) was performed using QuantiFast SYBR Green PCR Kit (Qiagen) and a StepOnePlus Real-Time PCR System (Applied Biosystems); the corresponding oligonucleotides are summarized in Supplementary Table 1. For each sample, Ct values were extrapolated using the StepOne software v2.3 and ratios of relative gene expression levels of *ATP5* (reference gene) and *GLI1* were calculated based on a modified ΔΔCt method taking into account different PCR kinetics^36^; PCR efficiencies are summarized in Supplementary Table 1.

### Confocal imaging, deconvolution and image analyses

The neuroepithelia of organoids were imaged with a Nikon A1R FLIM confocal microscope with the experimenter blinded to the genotype. Laser power and gain were adjusted to maximise intensity of the staining while avoiding overexposure. The Z-stack contained between 5μm and 15 μm of tissue section imaged in 0.13 μm steps. An optical zoom of x2.26 with pixel size of 0.06 was used to show more detail of the cilia. Deconvolution was performed using Huygenes Essential with the signal to noise ratio adjusted to values between 10 and 40 and the quality threshold set to 0.01.

Fluorescence mean intensity of ciliary markers relative to axonemal ARL13B staining were analysed using ImageJ software. 15 cilia per organoid (3 organoids per genotype) were chosen that had elongated neuroepithelia. For both, ARL13B and the marker of interest, background mean staining intensities were determined and deduced from the respective intensity levels in the cilium. The intensity ratio between the marker of interest and ARL13B was used for statistical analyses, thereby minimising bias that might have originated from a variability in the staining or image acquisition. For statistical analyses, the intensity ratios of all control and mutant organoids were collected in two separate groups. The length of primary cilia (15 cilia per organoid for 3 control and 3 mutant lines) was determined using ImageJ.

### Single-cell mRNA-seq and Bioinformatic Analyses

For each of the 3 control and 3 homozygous mutant lines, 10 D38 organoids were pooled, while 2 organoids for used for each line for the D94 single cell analysis. The organoids were minced into small pieces using a sterile razor blade and dissociated into single cell suspensions using a Worthington Papain Dissociation kit (Worthington Biochemical, #LK003150) as per manufacturer’s instruction, except for D94 organoids for which the Papain incubation time was increased to 75 minutes. After papain treatment, cells were centrifuged, resuspended in ice-cold 1x PBS and filtered using 40mm pluriStrainer Mini filter (Fisher Scientific, #431004050). The final cell concentration was adjusted to 1000 cells/ml in PBS.

Single-cell RNA-Seq libraries were prepared for the 10x scRNA sequencing platform according to the manufacturer’s instructions with the Chromium Next GEM Single Cell 3@ Reagent Kit v3.1 (Dual Index; 10x Genomics) with a target cell discovery of 10.000 cells. The average library size was 450bp. The resulting libraires were sequenced using Illumina NovaSeq. The single-cell sequencing reads were mapped to the human genome; per-cell, per-gene count matrices were produced using 10x CellRanger v.7.0.1^37^. Ambient RNA was estimated and removed using the SoupX R package v.1.6.2^38^. Doublets were identified and removed using the scDblFinder R package v.1.10.1^39^. Quality control, normalization and clustering of data were performed using the Seurat R package, v.4.1.0^40^. Cells expressing <250 or >5500 genes and/or 10% mitochondrial genes were excluded. Differentially expressed genes were identified for each cell group with the (i) FindMarkers() command in Seurat (Supplementary Tables 2 and 3), (ii) pseudo-bulk differential expression analysis (Supplementary Tables 4 and 5), performed by summarising single cell gene expression profiles at the subject level using the aggregateBioVar R package, version 1.8.0^41^, then calculating differentially expressed genes using DESeq2, version 1.38.3^42^. Pseudo-bulk analysis revealed few DEGs at FDR < 0.05. The FindMarkers() analysis was used to investigate potential patterns of gene expression changes in top genes, acknowledging that further work would be needed to definitively confirm individual gene differential expression. Gene ontology and network analyses were performed using clusterProfiler software^43^ in the annotation category BP (Supplementary Table 6). RNA velocity was estimated using scVelo^44,45^. The Seurat commands FindIntegrationAnchors() and IntegrateData() were used to integrate datasets with a publicly available dataset^46^. CellChat was employed to infer cell-cell communication^47^. Raw sequences were deposited in ENA (E-MTAB-15192).

### Quantification and statistical analysis

Data were analysed using GraphPadPrism 10 software with n=11-16 organoids for all analyses. Normal distribution was tested with Shapiro-Wilk or D’Agostino-Pearson omnibus normality tests and F-tests were used to test for equal variation. Normally distributed data with equal variance were analysed with unpaired t-tests but with unpaired t-tests with Welch’s correction if data showed unequal variance. In all other cases, Mann Whitney tests were used. For analyses containing more than three groups, one-way ANOVA tests were performed followed by multiple comparisons testing. A single asterisk indicates significance of p<0.05, two asterisks indicate significance of p<0.01, three asterisks of p<0.001 and four asterisks of p<0.0001. Graphs show the mean as well as upper and lower 95% confidence intervals. Statistical details can be found in the figure legends; Supplementary Table 7 provides a detailed summary of descriptive statistics of the tests used.

## RESULTS

### Generation and initial characterisation of *CEP41* mutant iPSCs and organoids

To determine the effects of *CEP41* ASD mutations on human cortical development, we first generated mutant human iPSC cell lines with a heterozygous or homozygous R242H mutation (Supplementary Figure 1) using a clustered regularly interspaced short palindromic repeats (CRISPR)/Cas9 approach. This missense mutation was identified in ASD individuals^25^ and was predicted to have a deleterious effect on the structure of CEP41’s evolutionarily conserved rhodamine domain^25^, that is presumed to act as a protein-protein interaction domain^27^. A guide RNA (gRNA) was selected that had at least three mismatches to potential off-target sites which are highly homologous to the on-target site. The gRNA/Cas9 plasmid was co-transfected with a template oligonucleotide carrying the R242H mutation into control iPSCs^48^. In this way, we identified 8 and 5 clones with the desired heterozygous and homozygous mutation, respectively, as confirmed by restriction fragment length polymorphism and Sanger sequencing (Supplementary Figure 1). Three homozygous mutant clones, three heterozygous and three wild-type clones that had not undergone gene editing were chosen for further analyses. All clones were karyotypically normal, except for one homozygous clone which showed a decreased telomeric dosage on the q arm of chromosome 22, and retained the pluripotency markers NANOG, OCT3/4 and TRA-1-60 (Supplementary Figure 2). In none of the clones did we detect off-target activity of the gRNA for the five highest candidate off-target sites as assessed by Sanger sequencing.

Control and *CEP41* mutant iPSC lines were differentiated into cerebral organoids using a modified Lancaster protocol (Supplementary Figure 1)^49^. After dual-Smad inhibition stimulating neural induction and embryoid body (EB) formation, FGF2 was added to the culture medium to promote neuroepithelial expansion. EBs were maintained on a shaking incubator to enhance oxygen exchange and nutrient absorption. By week 4, control and mutant organoids developed large neuroepithelial loops that continued to expand over the following two weeks, prior to harvesting the cerebral organoids at Day 38 (D38). Using these organoids, we started to determine the effects of the mutation on the CEP41 protein. Western blot and immunofluorescence analyses revealed that the R242H mutation did not affect CEP41’s stability nor its localisation at the basal body and at the ciliary axoneme (Supplementary Figure 1). Notably, CEP41 protein is absent from the mitotic spindle in control and mutant radial glial cells (RGCs) and spindle orientation is not affected by the *CEP41* mutation (Supplementary Fig. 3) suggesting that CEP41 is unlikely to play a role at the centrosomes during RGC mitosis. Taken together, these findings suggest that the R242H mutation does not affect CEP41’s stability and ciliary localisation.

### The *CEP41* mutation alters cortical progenitor development and the generation of glutamatergic projection neurons

To determine the effects of the *CEP41* R242H mutation on early corticogenesis, we used single-cell RNA sequencing (scRNA-seq) of D38 organoids. Accounting for clonal heterogeneity, we pooled 3 organoids from 3 control and 3 homozygous mutant lines. Uniform manifold approximation and projection (UMAP) and unbiased clustering of 42,776 cells revealed seven clusters of telencephalic cell populations: RGCs (RG1-5), basal intermediate progenitors (bIP1-2), excitatory neurons (EN1-3), ventral telencephalic progenitors (VP1-2), interneurons (IN1-3), Cajal Retzius cells (CR) and cortical hem/choroid plexus (HemCP) (Fig. 1A, B). These clusters expressed known marker genes and the presence of corresponding cells in the organoids was confirmed by immunofluorescence stainings (Fig. 1C-J). An 8^th^ cell group, termed RG+EN, contained RGCs as well as excitatory neurons and interneurons and was mainly composed of mutant cells that were characterised by high and intermediate expression levels of *LRRC75A* and several *histone* genes (*HIST1H3B*, *HIST1H3D*, *HIST1H2AG*), respectively (Supplementary Figure 4). Except for this cell group, however, all control and mutant lines contributed to all other cell clusters (Supplementary Figure 5).

**Figure 1:**
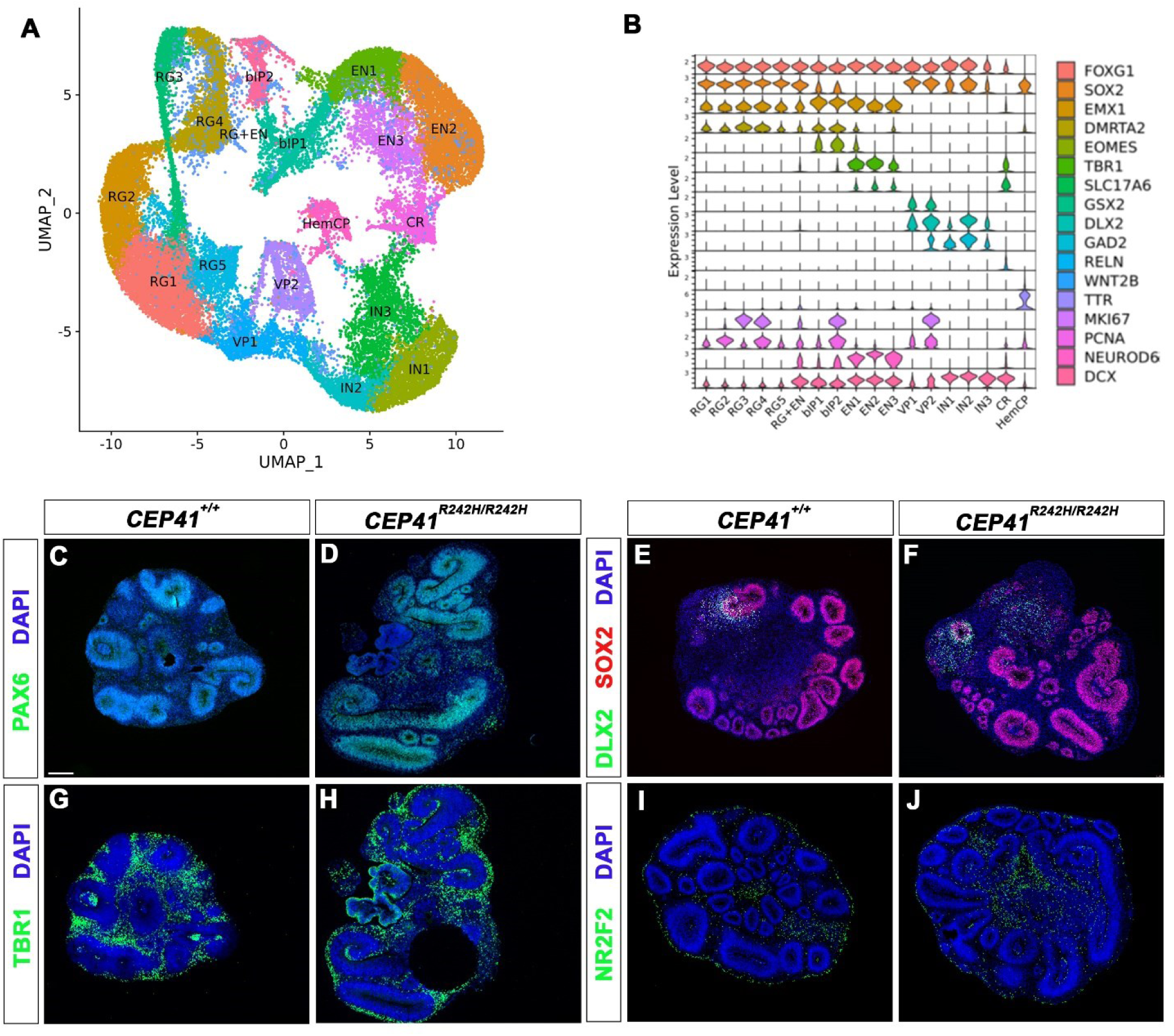
Expression of dorsal and ventral markers in *CEP41* mutant cortical organoids. (A) UMAP of the cortical organoid scRNA-seq data set revealing the presence of cortical progenitors and excitatory neurons, as well as the generation of interneuron progenitors and interneurons. (B) Violin plot indicating the markers used to identify different cell types in the single cell data set. (C-J) Immunofluorescence stainings with the indicated markers to confirm the presence of cortical progenitors (PAX6) (C, D), ventral progenitors (DLX2) (E, F), deep layer projection neurons (TBR1) (G, H) and interneurons (DLX2 and NR2F2) (E, F, I, J). Scale bar: 200μm.

We first focused our analysis on the dorsal telencephalic lineage. Determining the ratio between progenitor cell types and excitatory neurons we found an increased proportion of progenitors in mutant organoids driven by an increase in the number of RGCs (Fig. 2A). The RG cluster consisted of five subpopulations from which the RG1 and RG4 populations were predominately augmented (Fig. 2B). As the RG subpopulation mainly differed in the expression of cell cycle markers such as KI67 and PCNA, we assigned the cell cycle state to all individual RGCs and found that the fraction of cells in G2/M was increased at the expense of S phase cells (Fig. 2C). Indeed, acute BrdU labelling for 4 hours revealed a decline in the proportion of SOX2+ progenitors in S phase (Fig. 2D-I). Interestingly, while there were only a few gene expression changes in mutant RGCs (Supplementary Table 2-5), the RG2-4 populations presented with a significant ca. 1.3fold up-regulation of *HES5* which plays a role in maintaining the proliferative state of multipotent cortical progenitors^50,51^, consistent with the increased proportion of RGCs. Investigating the proportion of the three excitatory neuron population revealed a strong increase in the EN1 population at the expense of EN2 (Fig. 2J). Moreover, RNA velocity analysis showed that EN1 neurons likely represent newly born neurons whereas EN2 are more differentiated (Fig 2K, L). This idea is also supported by remnant expression of *EOMES* and low *NEUROD6* mRNA levels in EN1 (Fig. 1B). These findings suggest that the *CEP41* R242H mutation alters the balance between dorsal progenitor proliferation and differentiation and leads to a delayed formation of cortical excitatory neurons in D38 organoids.

**Figure 2:**
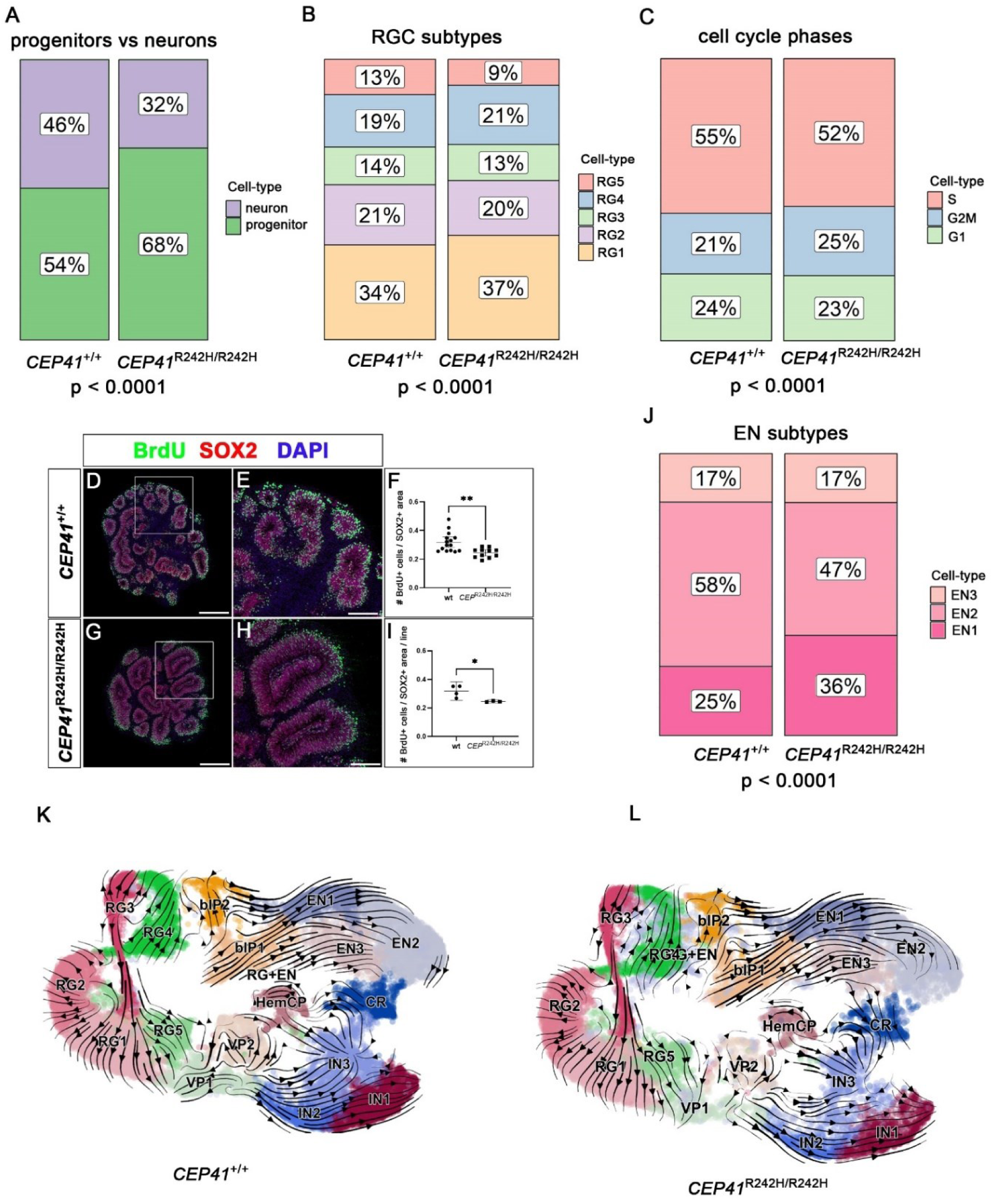
Altered cortical progenitor proportions in D38 *CEP41*^R242H/R242H^ mutant organoids. (A) Bar plots showing an increased ratio of cortical progenitors to excitatory neurons in mutant organoids. (B, C) Within the progenitor population, the proportions of RG1 and RG4 populations is augmented (B) reflecting a decrease in the proportion of S phase cells. (D-I) BrdU and SOX2 double labelling confirmed a reduced proportion of S phase cells. (J) Bar plot revealing an increased EN1 proportion at the expense of EN2. (K, L) RNA velocity analysis. Statistical data are presented as means ± 95% confidence intervals (CI). Chi-square tests (A-C, J); unpaired t-tests with Welch’s correction (n=4 for control and n=3 for mutant) (F, I); * p < 0.05. ** p < 0.01. Scale bars: 500 μm (D, G), 200 μm (E, H).

We next analysed the consequences of this imbalance at a later state of corticogenesis and repeated the single cell analysis with D94 organoids. Unbiased clustering led to the identification of apical and outer RGC (aRG and oRG) and several excitatory neuron clusters that were assigned as deep (EN(DL); EN(LIV-V); EN(mix)) and upper layer cortical neurons (EN(UL1-3)) based on marker gene expression (Fig. 3A-C). Organoids also contained a group of ventral progenitors (VP) and two interneuron populations (IN1-2). All these clusters contained cells from each the three control and mutant lines (Supplementary Figure 6). In contrast to the earlier stage, the proportion of excitatory neurons to dorsal progenitors was increased in D94 mutant organoids (Fig. 3D). Within the progenitor populations, mutant organoids showed an increased proportion of oRGs (Fig. 3E), a progenitor subtype predominantly generating late born upper-layer neurons^52^. Consistent with this idea, the investigation of neuronal subtypes showed an excess of upper layer neurons in mutants (Fig. 3F). Immunostainings with the upper and deep layer markers TBR1 and SATB2, respectively, confirmed this increase which was also observed in organoids heterozygous for the *CEP41* R242H mutation (Fig. 3G-J). Taken together, these findings suggest an early delay in neurogenesis which is followed by an increased formation of oRGs with a concomitant overrepresentation of upper layer neurons at D94.

**Figure 3:**
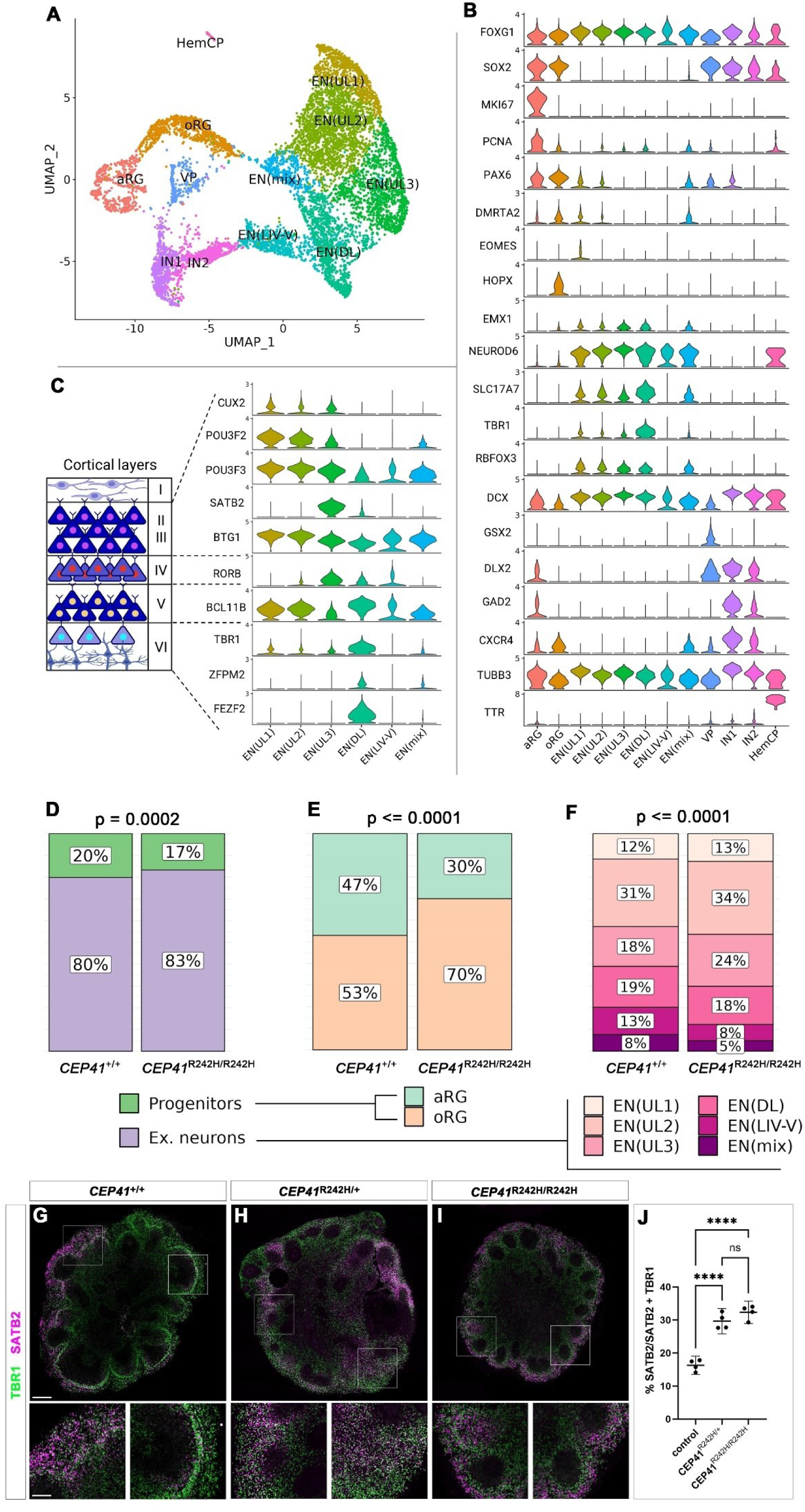
Increased oRG and upper layer cortical neuron formation in D94 *CEP41* mutant organoids. (A) UMAP showing 12 clusters of cell types identified in D94 cortical organoids. aRG/oRG, apical/outer radial glial cells; EN(UL1-3/DL/LIV-V/mix), excitatory neurons; VP, ventral progenitors; IN cortical interneurons; HemCP, choroid plexus. (B) Violin plot indicating the expressions of cell type-specific marker genes. (C) Violin plot indicating the expression of cortical layer-specific markers to distinguish excitatory neuron clusters. Upper layers (II-IV); deep layers (V-VI). (D-F) Bar plots showing altered cell-type proportions: dorsal progenitors (aRG and oRG combined) vs excitatory neurons (all ENs combined) (D), individual progenitor subtypes (E) and individual excitatory neuron subtypes (F). (G-I) Immunofluorescence stainings revealing the presence of deep (TBR1) and upper layer (SATB2) projection neurons. (J) Quantification revealing an increased percentage of upper layer cortical neurons in heterozygous and homozygous mutant organoids; n=4 lines for each genotype. Statistical data are presented as means ± 95% confidence intervals (CI). Chi-square tests (D-F). Yates’ continuity correction was applied to reduce the approximation error in D and E. One-way ANOVA with Holm-Sidak’s multiple comparisons test (J). **** p < 0.0001. Scale bars: 500 μm (G) and 100 μm in insets.

Since cilia are critical for cell signalling, we started to investigate potential signalling pathways that may lead to the increased formation of oRGs and upper layer cortical neurons using the CellChat tool^47^. We were particularly interested in pathways that were altered in late organoids but unchanged in earlier ones. This analysis showed an up-regulation of Pleiotrophin (PTN)/midkine signalling specifically in all progenitor populations of D94 organoids (Supplementary Figure 7). Its receptor, PTPRZ1, is widely expressed in apical and outer RGCs^53^. Interestingly, *PTN* expression is significantly increased in mutant ventral and dorsal aRGs and in oRGs (Supplementary Figure 7) suggesting that this up-regulation may lead to increased proliferation of oRGs and a subsequent augmented formation of upper layer cortical neurons.

### Interneuron development is affected in *CEP41* mutant organoids

The next set of experiments examined the impact of the *CEP41* mutation on the ventral telencephalic lineage. We first employed the single cell data sets to examine whether the proportion of ventral telencephalic cell types was altered as for the dorsal lineage. In D38 organoids, there was no change in the proportions between the two progenitor cell types, but a slight increase in interneurons relative to progenitors (Supplementary Figure 8). Within the interneuron populations, IN1 and IN3 were increased at the expense of IN2. In contrast, there was a remarkable decline in the numbers of ventral cell types in D94 organoids with ventral progenitor and interneuron cell clusters largely consisting of control cells (Supplementary Figure 8). Nevertheless, the proportion of progenitors and interneurons was not affected while IN1_D94 interneurons were significantly underrepresented within the interneuron population. These findings suggest that the *CEP41*^R242H/R242H^ mutation affects the differentiation and survival of cortical interneurons. To gain insights into the identity of these interneuron clusters, we integrated our data set with published single cell data derived from human foetal, primary ventral telencephalic tissue^46^ (Supplementary Figure 8). This integration revealed that the IN1 cell cluster largely consisted of lateral ganglionic eminence (LGE) derived striatal interneurons. IN2 and IN1_D94 cells mainly co-localized with interneurons derived from the caudal ganglionic eminence (CGE) but IN2 also made a significant contribution to medial ganglionic eminence (MGE) interneurons. Finally, IN3 and IN2_D94 cells formed a separate cluster but contributed to all three combined GE clusters. These patterns were confirmed by the expression of markers specific for each of the ganglionic eminences (Supplementary Figure 9).

We next sought to determine whether changes in cell proportions coincided with altered gene expression in each of the ventral telencephalic cell populations. Investigating patterns of gene expression changes in top genes, we noted a larger number of differentially expressed genes (DEGs) in D38 ventral cell clusters than in dorsal RGCs and excitatory neurons (Fig. 4, Supplementary Table 2-5). Gene ontology (GO) analysis revealed that up-regulated genes in VP1/2 and IN1-3 cell clusters of D38 organoids were primarily involved in protein formation and maturation (GO:BP terms: “protein targeting to ER”, “SRP dependent co-translational targeting to membranes”, “ribosome biogenesis”) (Fig. 4A, B, Supplementary Figure 10). In fact, most of the up-regulated genes in the IN3 cluster encode proteins of the small and large ribosomal subunits. The role of down-regulated genes varied more between the different cell groups. These genes were associated with splicing in VP1 progenitors, whereas genes involved in NADH generation and glycolysis (*ENO1/2*, *PKG1*) were found to be differentially expressed in the VP2 cluster (Fig. 4C, D; Supplementary Figure 10). Moreover, the *CEP41*^R242H/R242H^ mutation affected processes involved in neuronal differentiation in all three populations of interneurons, emphasized by GO:BP terms such as “axonogenesis”, “neuron migration”, “GABAergic interneuron differentiation” and “postsynaptic cytoskeleton organization” suggesting a delay in neuronal differentiation. Gene set enrichment analysis (GSEA) also indicated an increase in Myc target gene expression, but a reduction in glycolysis, hypoxia and mTORC1 signalling (Fig. 4E).

**Figure 4:**
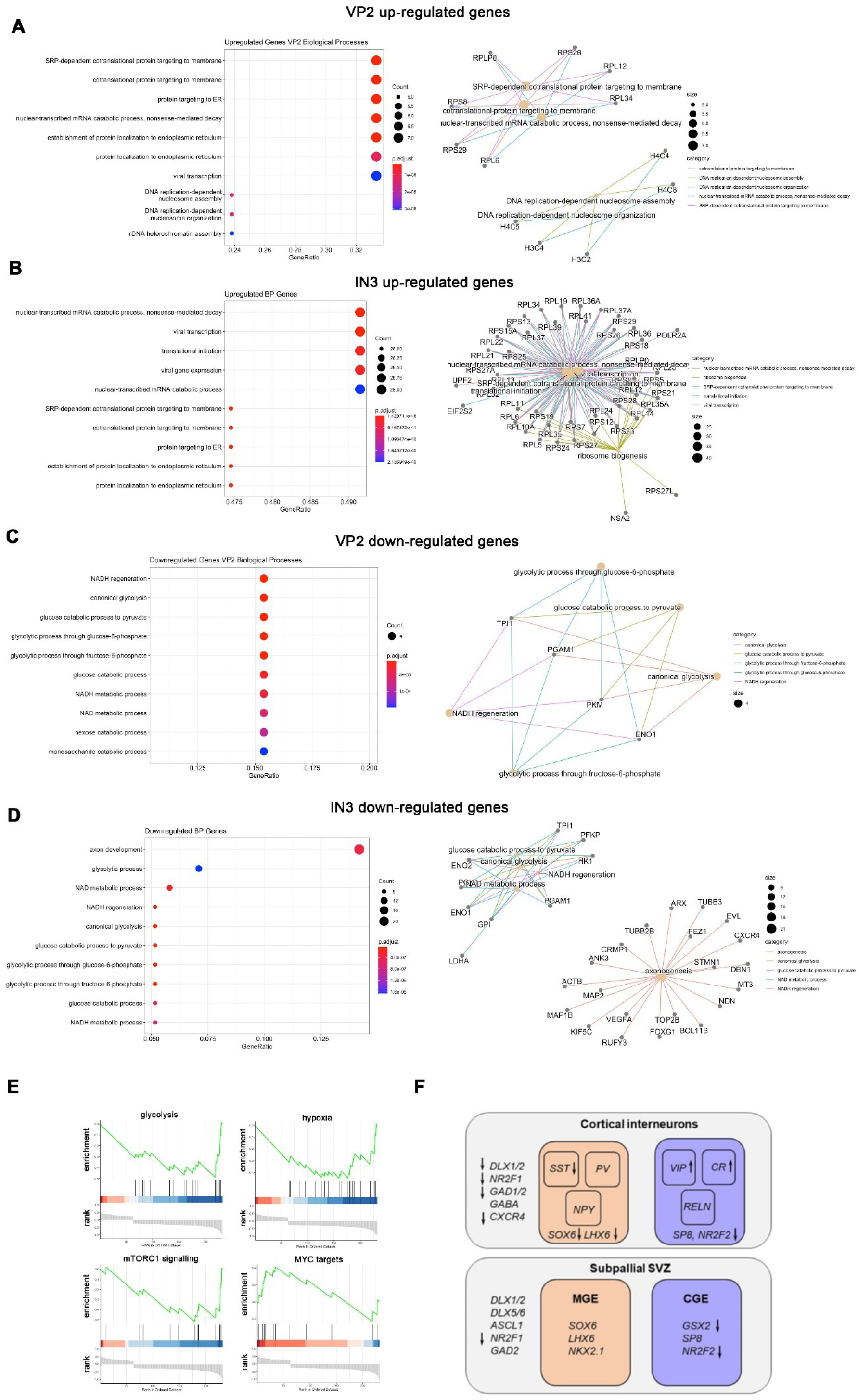
Analysis of gene expression changes in D38 ventral cell types. (A-D) Gene ontology (GO) and network analyses of up-regulated (A, B) and down-regulated genes (C, D) in VP2 (A, C) and IN3 (B, D). GO terms are ordered by increasing *p*-value from top to bottom; *p*-values are shown colour coded. Network analysis showing DEGs associated with the GO terms. Only significant DEGs and GO terms (adjusted *p*-value<0.05) were considered. (E) Gene set enrichment analysis (GSEA) of glycolysis, hypoxia, mTORC1 signalling and Myc target gene expression. (F) Schematic illustrating differential expression of key regulatory genes as indicated by arrows in MGE and CGE progenitors and in cortical interneurons derived from these progenitors.

In addition to these altered processes, we noted that the expression of several key regulators of interneuron development were affected by the *CEP41*^R242H/R242H^ mutation (Fig. 4F). These include transcription factors that are essential for establishing and maintaining a general interneuron cell fate (*DLX1/2*, *NR2F1*) or specifically act in the MGE (*SOX6*, *LHX6*) or CGE lineages (*GSX2*, *NR2F2*). Most notably, *NR2F1* and *NR2F2* were amongst the strongest down-regulated genes in all progenitor and interneuron cell groups in D38 organoids. In addition, the expression of genes encoding GABA producing enzymes (*GAD1/2*) were affected as well as genes important for interneuron migration (*CXCR4*, *NR2F1/2*) or characteristic of interneuron subtypes (*SST*, *VIP*, *CR*). Taken together, these findings suggest that the *CEP41*^R242H/R242H^ mutation severely impacted on interneuron development in D38 organoids.

The analysis of D94 ventral progenitors revealed numerous up-regulated (n=91) and down-regulated (n=12) DEGs (Supplementary Figure 11). Interestingly, the GO terms for up-regulated DEGs were associated with axon and synapse development, processes active in neurons rather than progenitors. Since all down-regulated DEGs encoded ribosomal proteins, their GO terms described ribosomal activity. These findings suggest a premature expression of neuronal genes and down-regulation of ribosomal genes unlike the earlier stage. Moreover, interneurons showed few significant DEGs, likely due to the small sample size. Nevertheless, it is worth mentioning that IN1 showed a down-regulation of *NR2F1* and *EPHA5*, which both promote interneuron specification and migration^54,55^. These results suggest altered expressions of important developmental regulators in D94 *CEP41*^R242H/R242H^ ventral progenitors and interneurons.

To further characterize the DEGs and to gain insights into their association with ASD, we performed pathway analysis with the Simons Foundation Autism Research Initiative (SFARI) repository^56^. The progenitor and interneuron DEGs from both ages showed statistically significant overlaps with the set of ASD-linked genes listed in the SFARI database (Fig. 5A, B). Further comparison revealed that the overlap at D38 mainly stemmed from SFARI genes in the syndromic category with IN3 being the most affected population (Fig.5C). At D94, there were overlaps in all three gene score categories mainly with genes differentially expressed in the VP cluster. Enriched GO terms for the sets of overlapping DEGs included “negative regulation of neuron differentiation” and “forebrain development” in the younger organoids and “regulation of ion transmembrane transport” and “membrane depolarisation” in the older organoids (Fig. 5D-G). Moreover, overlapping genes are highly interconnected. These results strongly suggested that the gene expression changes caused by the *CEP41*^R242H/R242H^ mutation are highly relevant to ASD aetiology.

**Figure 5:**
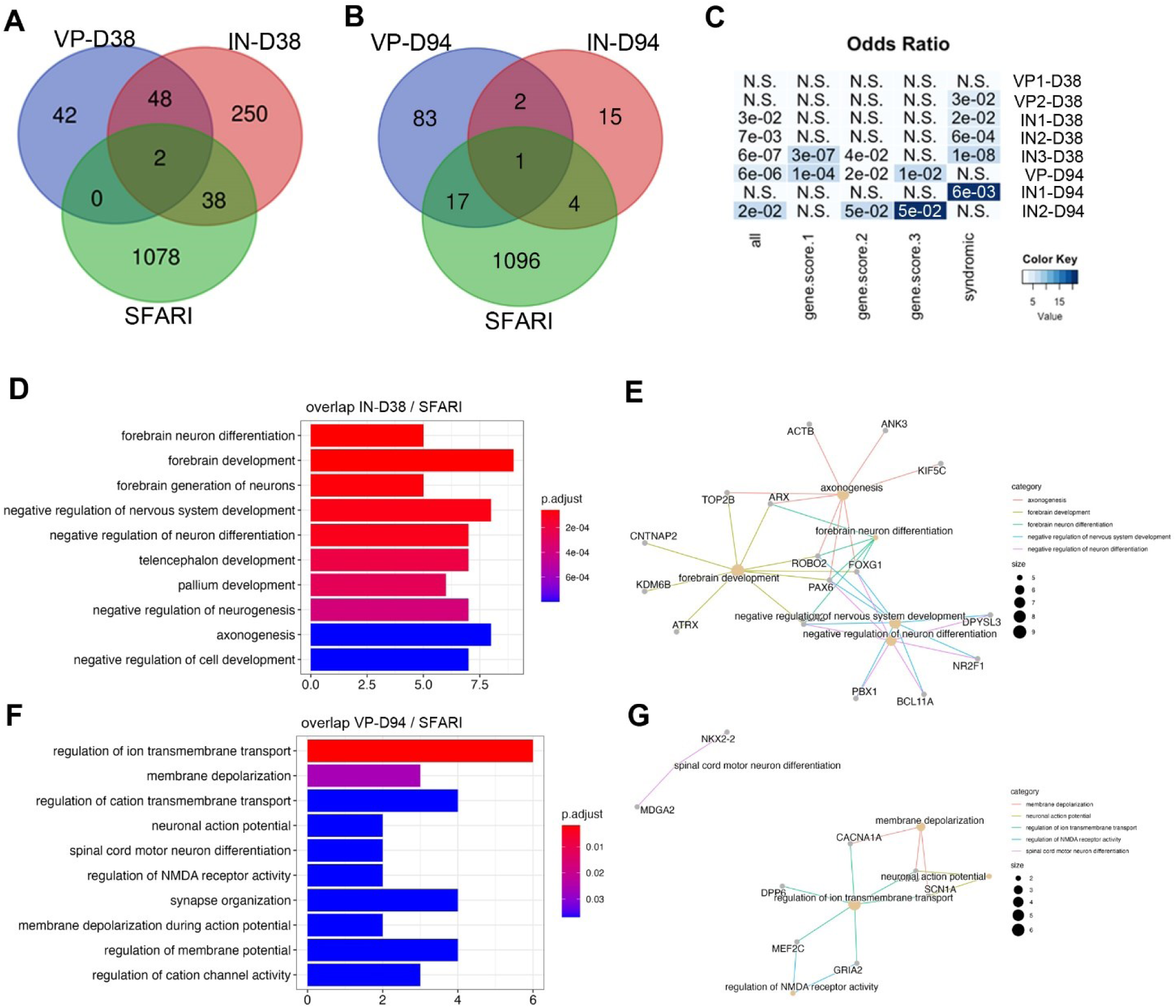
Gene expression changes in mutant interneurons and their progenitors are associated with ASD. (A-B) Venn diagrams intersection of differentially expressed genes in progenitors (VP; blue), interneurons (IN; red) and the SFARI list of ASD candidate genes (green) at D38 (A) and D94 (B). (C) Significance and odds ratio for the overlap between the various progenitor/interneuron populations and either the complete set or subcategories of ASD candidate genes. (D-G) GO analysis (D, F) and network plot (E, G) of the gene overlap between ASD candidate genes and IN-D38 (D, E) or VP-D94 (F, G).

### Mechanisms underlying altered interneuron development in *CEP41* mutant organoids

We next investigated the mechanisms of altered interneuron development in *CEP41* mutant organoids. Given its prominent role in ventral telencephalic development, we first investigated SHH signalling. We first interrogated the scRNA-seq datasets for the expression of genes involved in SHH signalling (*SHH*, *PTCH1*, *PTCH2*, *SUFU*, *GLI2*, *GLI3*) or of known target genes (*NKX2.1*, *OLIG2*, *HHIP*, *GLI1*) relative to the expression of the housekeeping genes *ACTB*, *ATP5F1D*, and *GAPDH*. While the latter were expressed abundantly, SHH signalling markers were only detected in a few cells, probably due limited sequencing depths, making this analysis inconclusive (Supplementary Figure 12). To overcome these limitations, we generated ventral telencephalic organoids^49,57^ which were immunostained for various ventral-specific markers to validate their ventral identity and analyse cell population sizes. For each immunostaining, the number of marker-expressing cells was scaled by the DAPI+ area to account for organoid size. Staining for the progenitor marker SOX2 revealed no differences in SOX2+ cell density in mutants (Supplementary Figure 13). Organoids were next stained for DLX2 and OLIG2, which are strongly expressed in MGE and LGE progenitors^58–60^, but neither DLX2+ nor OLIG2+ cell densities were affected in the mutant. NR2F2 is highly expressed in CGE-derived progenitors and interneurons^58^. Accordingly, we detected SOX2+/NR2F2+ double-positive and SOX2-/NR2F2+ single-positive cells in the organoids. Consistent with the *NR2F2* down-regulation in cortical organoids, we observed a significant, approximately 2-fold decrease in the NR2F2+ cell density in mutant ventral telencephalic organoids. Taken together, these findings confirm the ventral identity of *CEP41*^R242H/R242H^ organoids and indicate a perturbed development of NR2F2+ CGE-derived interneurons consistent with the scRNA-seq analysis.

We next employed *CEP41*^R242H/R242H^ ventral organoids for analysing SHH signalling. Activity of the SHH pathway was examined using qRT-PCR for the SHH target gene *GLI1*. Mutant organoids displayed an approximately 2-fold, marginally non-significant decrease of *GLI1* expression (Supplementary Figure 12). These results suggest a trend for decreased SHH signalling in *CEP41*^R242H/R242H^ ventral telencephalic organoids.

Finally, we aimed to explore a potential link between CEP41 and cilia on the one hand and the altered transcription factor network on the other. Recent perturbation experiments in human cortical organoids highlighted a requirement of *GLI3* for establishing cortical fate and controlling LGE vs MGE development^61^. Inspecting a published GLI3 cut & tag data set^61^ detected GLI3 binding to the *DLX1/2* and *DLX5/6* genes with extended binding regions near the *NR2F1/2* genes (Supplementary Figure 14). Given the widespread *NR2F1/2* dysregulation in *CEP41* mutant ventral progenitor and interneuron populations, we examined a human cortical organoid NR2F1 ChIP-seq data set^62^. This analysis demonstrated strong NR2F1 binding to its own promoter as well as to *LHX6*, the *DLX* genes, *GAD1* and *CALB1* (Supplementary Figure 15). Interestingly, this binding was evolutionarily conserved between human and mouse^63^.

### The R242 mutation affects tubulin polyglutamylation and IFT transport

To gain insights into the ciliary defects that may underlie the defective development of projection neurons and interneurons, we analysed the expression and localisation of several cilia markers. CEP41 has a prominent role in the ciliary transport of TTLL6^27^ that catalyses tubulin polyglutamylation, a modification critical for the stability of ciliary microtubules^64–66^. Consistent with this known CEP41 function, ciliary tubulin polyglutamylation levels were reduced in both, heterozygous and homozygous mutant RGCs (Fig. 6A-D). This reduced expression coincided with a shortening of cilia (Fig. 6E). Hypoglutamylation was shown to lead to a decrease in anterograde transport but not in retrograde transport^66^. Accordingly, expression levels of the anterograde transport component IFT88 were not only reduced in the axoneme, but IFT88 expression was also absent from the ciliary tip of *CEP41* homozygous mutant cilia (Fig. 6F-L). In contrast, ciliary expression and localisation of the retrograde transport component IFT144 was not affected (Fig. 6M-S). Finally, we noted that the expression of the gene encoding Phospolipase γ2 (PLCγ2) that acts downstream of PDGF signalling to induce deciliation^67^ was reduced in most D38 scRNA-seq cell clusters (Supplementary Tables 2 and 4). Taken together, these findings suggests that the R242H mutation affects cilia stability and might thereby reduce the signalling capacity of the cilium.

**Figure 6:**
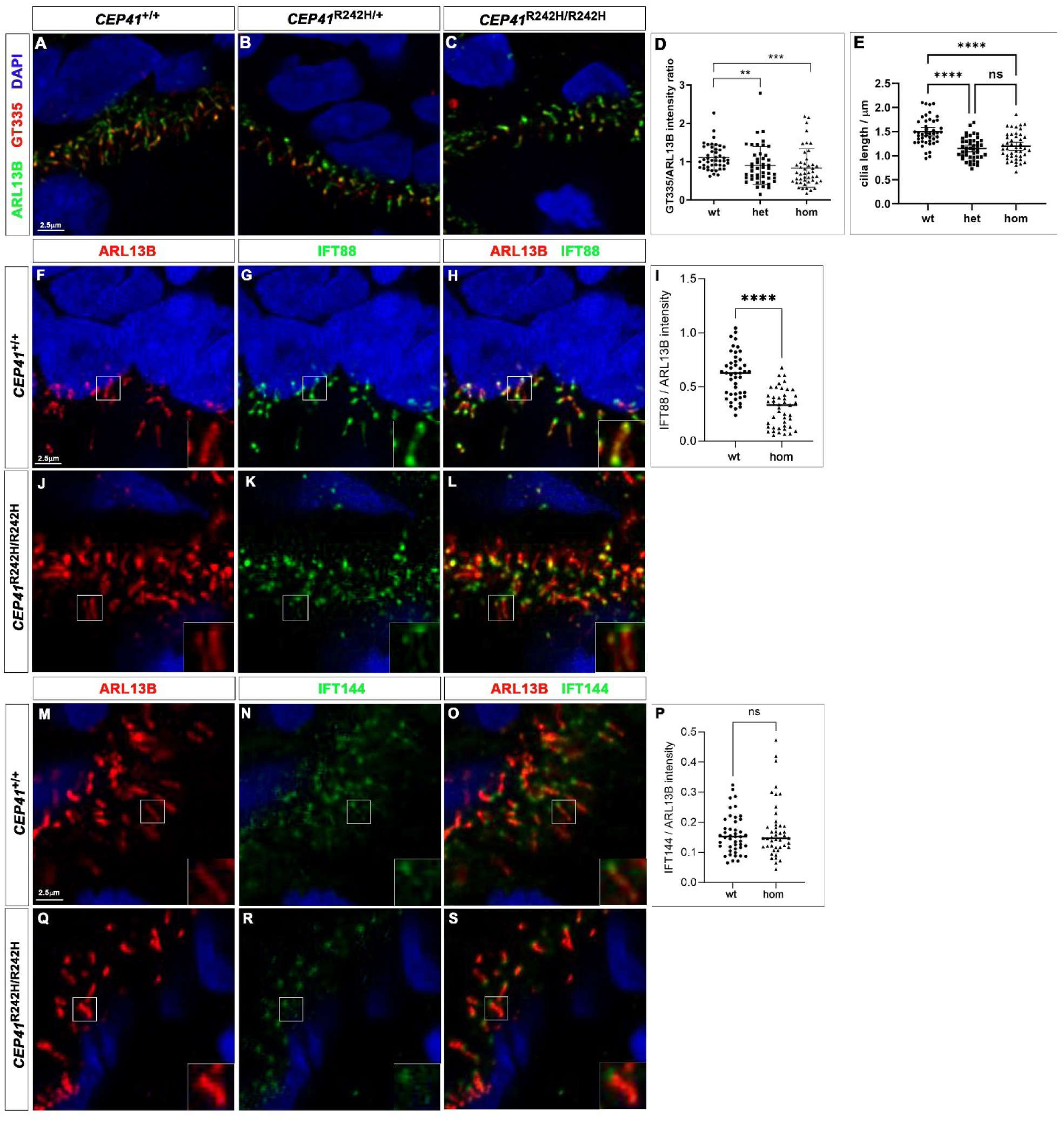
Ciliary alterations in *CEP41*^R242H/R242H^ organoids. Control and *CEP41*^R242H/R242H^ organoids were immunostained with the indicated markers. (A-E) Tubulin polyglutamylation (A-D) and ciliary length (E) are significantly reduced in heterozygous and homozygous mutant organoids. (F-S) Reduced IFT88 (F-L) but not IFT144 (M-S) expression in homozygous mutant organoids. Statistical data are presented as means ± 95% confidence intervals (CI); Kruskal-Wallis test followed by Dunn’s multiple comparisons test (D), one-way ANOVA followed by Tukey’s multiple comparisons test (E), unpaired t-test (I) and Mann Whitney tests (P); n=3 for control and mutant lines. n = 45 cilia for all three genotypes from three different lines; ** p < 0.01; *** p < 0.001; **** p<0.0001. Scale bar: 2.5 μm.

## DISCUSSION

This study introduces a framework to explore the molecular and cellular foundations of primary cilia and their involvement in the aetiology of ASD within a human context. Utilizing cortical and ventral telencephalic organoid models, our research showed that a *CEP41* ASD mutation impairs the development of both excitatory neurons as well as inhibitory interneurons. A delay in early neuronal differentiation led to an increased proportion of upper layer cortical neurons. Interneurons and their progenitors showed gene expression alterations, particularly affecting key developmental regulators and ASD candidate genes. These results indicate that the *CEP41* mutation might disrupt the E/I balance and thereby contribute to ASD pathogenesis.

### Effects of the *CEP41* mutation on cilia structure and signalling

Increasing evidence implicate altered corticogenesis during the second and third trimester of foetal development as a major pathomechanism in a large proportion of ASD subjects^8,9^. During this period, cell-cell signalling plays a dominant role and, indeed, recent studies highlighted an emerging role for primary cilia in the aetiology of NDDs and ASD in particular. Many ciliopathy patients presenting neurological symptoms are also diagnosed with ID and ASD^13^ while mutations in several high confidence ASD candidate genes also impact on cilia structure and/or signalling^14–19^. Interestingly, defective Shh signalling and dendritic arborisation were rescued by reinstating cilia in primary cultures of mouse cortical neurons mutant for *Mecp2*^16^ but in all other cases it remains to be seen whether these ciliary defects contribute to pathogenesis. Finally, mutations in various ciliary genes have been identified in ASD individuals, impacting ciliary gene transcription (*RFX3/4/7*), ciliogenesis (*FAM92B*, *OFD1*, *PCM1*), the transition zone (*AHI1*, *CEP290*), ciliary transport via the BBSome (*BBS4*) and the axonemal protein CEP41. Taken together, these findings suggest that ciliary dysfunctions contribute significantly to the pathogenesis of neurodevelopmental disorders in general and ASD in particular. However, our insights into roles of cilia in neural development in health and disease are mainly derived from animal or cell culture models, but differences in brain development between human and non-human mammals^68–70^ make extrapolating these results to human disorders difficult. Addressing this knowledge gap and given the proposed role of an E/I imbalance, we made use of human cortical and ventral telencephalic organoids to investigate the effects of an ASD mutation in a ciliary gene on excitatory and inhibitory neuron development. Amongst the ASD mutant ciliary genes, *CEP41* stood out with over 20 familial and sporadic ASD cases^23,25^. The affected individuals carry missense mutations that mainly cluster in the rhodamine domain. Based on structure predictions and analysis of zebrafish neurogenesis, we chose to investigate the R242H mutation, noted for its deleterious effect^25^. While ASD subjects carry a heterozygous mutation, our study primarily used homozygous mutant organoids as a proof of principle, with key findings replicated in heterozygous mutant organoids. This mutation did not affect the stability of the CEP41 protein nor its localisation at the basal body and the axoneme. Although centrosomal proteins can serve multiple functions in the cilium and during mitosis^71^, CEP41 protein did not localize to the mitotic spindle nor was spindle orientation affected in mutant RGCs, suggesting that altered organoid development is due to impaired ciliary signalling. Previous experiments in fibroblasts derived from JS subjects showed that *CEP41* is essential for transporting TTLL6 into the cilium for tubulin polyglutamylation^27,72^, while *Cep41* knock-down in cultured murine cells resulted in tubulin hypoglutamylation^27,65^. This posttranslational modification controls SHH signalling as well as ciliary stability and length^64–66^, a modifier of the cilia’s ability to sense and transduce extracellular signals^73^. Accordingly, we observed reduced levels of tubulin glutamylation in *CEP41*^R242H^ heterozygous as well as homozygous mutant RGCs suggesting a potential hypomorph or dominant negative effect. Cilia were also shorter with reduced expression of IFT88 but not IFT144 along the axoneme and at the ciliary tip. Altered IFT transport was recently identified as a consequence of tubulin hypoglutamylation and led to reduced activation of Shh signalling, probably due to a prolonged entry of Smo and Gli3 into the cilium^66^. *Cep41* knock-down in NIH 3T3 fibroblasts also impaired ciliary tip localisation of Gli2 and Gli3 upon activation of Shh signalling^65^. These findings help to explain the tendency for reduced SHH signalling in ventral telencephalic *CEP41*^R242H/R242H^ mutant organoids. Finally, *Cep41* was implicated in ciliary disassembly via AURKA activation^72^. This control mechanism might act in combination with the reduced expression of PLCγ2 which is crucial for inducing deciliation upon activated PDGF signalling^67^. Thus, the combined roles of CEP41 on ciliary stability and disassembly suggest that the *CEP41* ASD mutation might result in cilia with compromised stability and signalling capacity.

### Altered development of excitatory projection neurons and inhibitory interneurons and its relevance to ASD

Our study elucidates several potential mechanisms by which the *CEP41* mutation could contribute to the development of autism at the cellular level. A prevailing hypothesis postulates that an imbalance between excitation and inhibition plays a critical role in NDD aetiology. Although research on NDDs has primarily concentrated on changes in neuronal connectivity and circuitry, growing evidence also implicate defects occurring during foetal development of projection neurons and interneurons^8,9^. Interestingly, *CEP41* mutant organoids show alterations in developing excitatory and inhibitory cell lineages. Although gene expression changes were minimal in dorsal cell types, progenitor proliferation and differentiation appeared disrupted. An increased proportion of progenitors suggested delayed neuron formation in early organoids resulting in an increased proportion of oRGs and upper layer neurons at later stages. Differential gene expression identified increased *PTN* expression as a potential cause for the overproduction. *PTN* encodes a secreted, extracellular matrix associated growth factor linked to controlling oRG proliferative behaviour^53^ and stimulates the epithelial to mesenchymal transition in glioblastoma cell lines^74^ which has been associated with the rapid migratory bursts preceding oRG cell division^75^. The elevated proliferation and overproduction of cortical neurons align with reports of excess cortical neurons ^76^, increased brain weights^76–78^ and early brain overgrowth in ASD^8,79–82^. In mice, excess prenatal neurogenesis similarly caused an overabundance of upper-layer cortical neurons, an imbalance of excitation and inhibition, altered neural functioning, and ASD-like behaviours^83^.

Our study also highlights significant disruptions in cortical interneuron development. Unlike the excitatory dorsal cell lineage, ventral telencephalic cell populations displayed notable gene expression changes. Key among these are the *NR2F1/2* genes, known to control cortical arealisation and excitatory neuron as well as interneuron differentiation in mice^55,84–90^. These were differentially expressed in all ventral telencephalic populations, with recent genomic analyses revealed *NR2F1* mutations in ASD patients ^91–94^. Modelling one of these mutations in human cortical organoids led to increased interneuron formation due to elevated SHH signalling^62^. Moreover, the same mutation caused an E/I imbalance and behavioural ASD-like deficits in mice^62^, further underscoring *NR2F1*’s critical role.

In addition, the mRNA levels of several other key transcriptional regulators of interneuron development, including *ARX*, *DLX2*, *GSX2*, *LHX6* and *SOX6*, were altered. Although these changes were generally milder and confined to a subset of ventral telencephalic populations, they may have implications for ASD aetiology. The expression of *Dlx1/2* and *Dlx5/6* is controlled by internal enhancers. Their deletion caused a twofold reduction in *Dlx* gene expression, leading to behavioural abnormalities in mice^95^. Intriguingly, mutations were also unveiled in the *DLX5/6* internal enhancer in individuals with autism^96,97^. Moreover, *Dlx* genes form a transcriptional circuitry with other transcription factors^63^, many of which were differentially expressed in mutant organoids. For example, *DLX1/2* directly controls the expression of the aristaless-related homeobox gene *ARX*^98^, a gene linked to various neurodevelopmental disorders^68,99^. Finally, differentially expressed genes were significantly overrepresented in the SFARI ASD database. This overlap combined with GO/gene network analyses showing a potential disruption of early neuron formation, differentiation and synapse development underscores the impact of the *CEP41* mutation on interneuron development in our organoid model.

Additionally, there was a marked up-regulation of genes encoding ribosomal proteins, particularly in the IN3 population in D38 organoids, which exhibited an altered expression of 40 out of 85 ribosomal proteins. Altered synaptic protein translation has been highly implicated in ASD aetiology^100^. While it remains to be elucidated whether synaptic ribosome composition and function are altered in *CEP41* mutant interneurons, other translation-based mechanisms may influence interneuron differentiation. During development, ribosome biogenesis and global protein synthesis rates are not uniform but tightly and dynamically regulated by Myc dependent transcription of genes encoding ribosomal proteins and by mTORC1 signalling, respectively^101–104^. Global protein synthesis rates are consistently lower in stem cells including neural stem cells, increase during early differentiation and are reduced again in later differentiation. In turn, ribosome biogenesis is selectively high in stem cells, drops at early differentiation stages and rises again at later differentiation. An up-regulation of myc target genes and a down-regulation of mTORC1 signalling in IN3 interneurons, as revealed by GSEA, therefore suggests that these cells might be an immature, plastic state which is consistent with the IN3 contribution to CGE, LGE and MGE interneurons. Taken together with the strong overlap of IN3 DEGs and the SFARI list, this finding suggests that the *CEP41* ASD mutation might affect interneuron differentiation at an early developmental stage. In addition, altered ribosomal protein expression can also affect ribosome heterogeneity, an important developmental regulator. Translational control was also found to modify cell signalling and the SHH pathway in particular. The *Ptch1* 5’ untranslated region contains an eIF3c binding site and several upstream open reading frames (ORFs) competing with the main ORF for ribosomes. These elements positively and negatively regulate Ptch1 protein translation, respectively, during limb and neural tube patterning^105,106^, raising the intriguing possibility that changes in ribosomal protein expression might influence SHH signalling and thereby differentiation of *CEP41* mutant interneurons.

These interneuron lineage specific gene expression changes may contribute to ASD behaviours in the *CEP41* patients and are consistent with observations on the role of interneurons in ASD. Interneurons modulate neural circuits, generate cortical oscillations and maintain the E/I balance^68^, all of which can be disrupted by changes in interneuron numbers or inhibitory synapses^5,107^. In turn, perturbing high-confidence ASD-risk genes in human cortical organoids and in mouse telencephalic interneurons increased the proportion of ventral cells and led to repetitive and restricted behaviours, respectively^108–111^. Taken together with the altered proportions of excitatory neurons, these findings suggest that the *CEP41*^R242H/R242H^ mutation induces an E/I imbalance, a hypothesis that warrants further investigations.

## ACKNOWLEDGEMENTS

We are grateful to Drs Thomas Becker, Calvin Chan, John Mason and David Price for critical comments on the manuscript and Dr Tilo Kunath for sharing the NAS2 line. This work was supported by a grant from the Simons Initiative for the Developing Brain (SFARI -529085) to TT. NCH is supported by a Wellcome Trust Senior Research Fellowship in Clinical Science (ref. 219542/Z/19/Z).

## CONFLICT OF INTEREST

The authors declare no competing financial interests in relation to the work described.

## SUPPLEMENTARY FIGURES

**Supplementary Figure 1:**
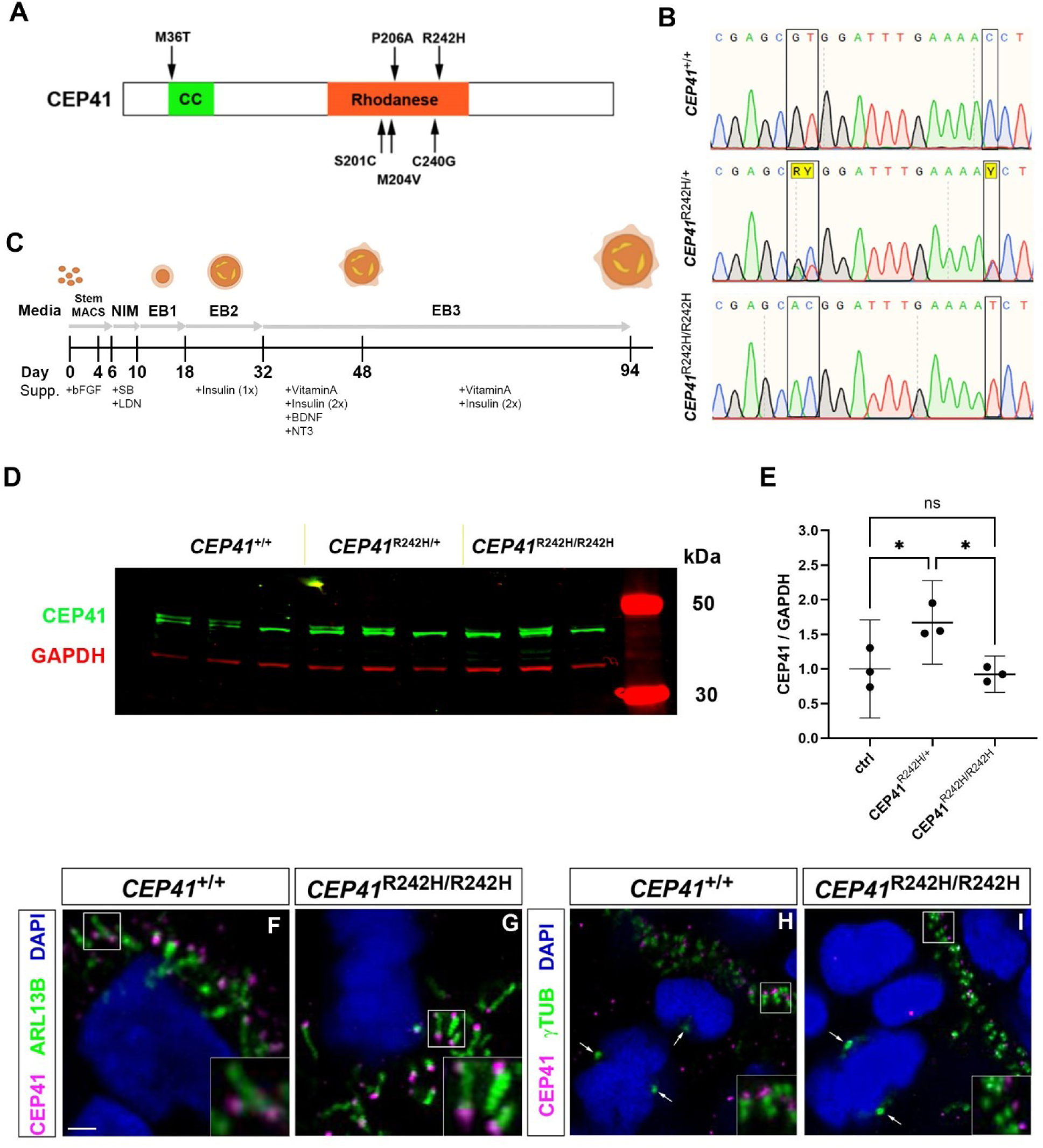
*CEP41* mutagenesis and CEP41 protein expression in cortical organoids. (A) Schematic domain structure of the CEP41 protein with indication of missense mutations found in ASD individuals. (B) Sequencing traces confirming successful mutagenesis. In addition to the R242H mutation, a second silent mutation was introduced to inactivate the PAM site. (C) Schematic of the protocol used to generate cortical organoids. (D) CEP41 Western with protein extracts from D38 control, heterozygous and homozygous mutant organoids. (E) Quantification of the CEP41 Western blot. (F, G) CEP41 expression in cilia of control (F) and in *CEP41*^R242H/R242H^ organoids (G). CEP41 protein was confined to the basal body with lower expression in the axoneme in both genotypes. (H, I) CEP41 protein is absent from the mitotic spindle as indicated by double staining for γTubulin (arrows). Statistical data are presented as means ± 95% confidence intervals (CI); one-way ANOVA followed by Tukey’s multiple comparisons test with n=3 for all genotypes (E). * p < 0.05. Scale bar: 2.5 μm.

**Supplementary Figure 2:**
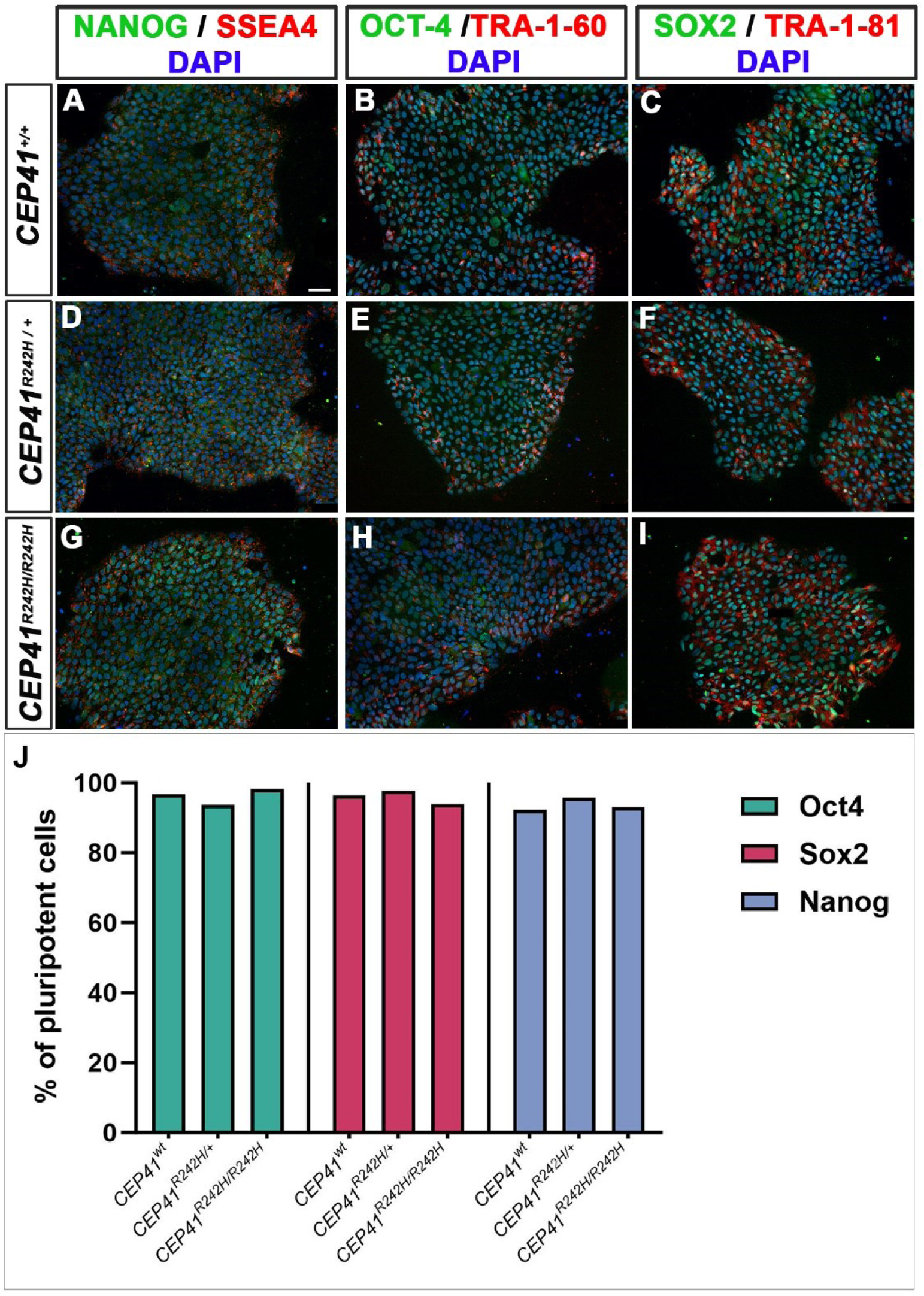
Expression of pluripotency markers in control, *CEP41*^R242H/+^ and *CEP41*^R242H/R242H^ iPSC lines. (A-I) Immunofluorescence stainings for the indicated markers. All iPSC lines were positive for NANOG (A, D, G), OCT4 (B, E, H), and SOX2 (C, F, I). (J) Schematic diagram depicting the percentage of cells positive for the indicated stem cell markers. Scale bar: 50 μm.

**Supplementary Figure 3:**
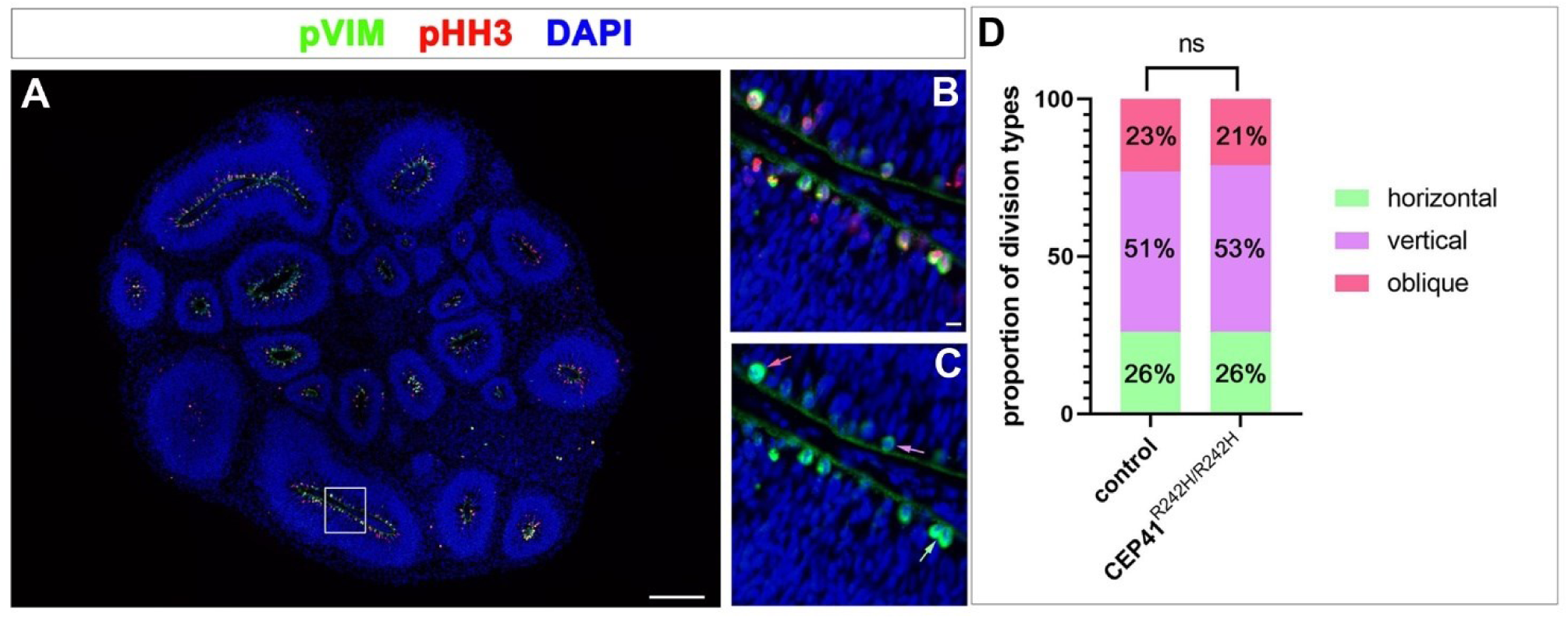
Orientation of the mitotic spindle is not altered in *CEP41* mutant apical radial glial cells. (A-C) Immunofluorescence staining against pHH3 and pVIM to reveal mitotic cells and the orientation of the mitotic spindle, respectively. The white arrow indicates a horizontal, the purple arrow a vertical and the red arrow an oblique cell division plane (C). (D) Quantification of horizontal (n=238 and 319 mitoses for control and homozygous mutant organoids, respectively, derived from three different lines for each genotype), vertical (n=469 and 647 mitoses for control and homozygous mutant organoids, respectively, derived from three different lines for each genotype) and oblique (n=214 and 257 mitoses for control and homozygous mutant organoids, respectively, derived from three different lines for each genotype) cell divisions. Chi-square test; p=0.9374. Scale bars: 200μm.

**Supplementary Figure 4:**
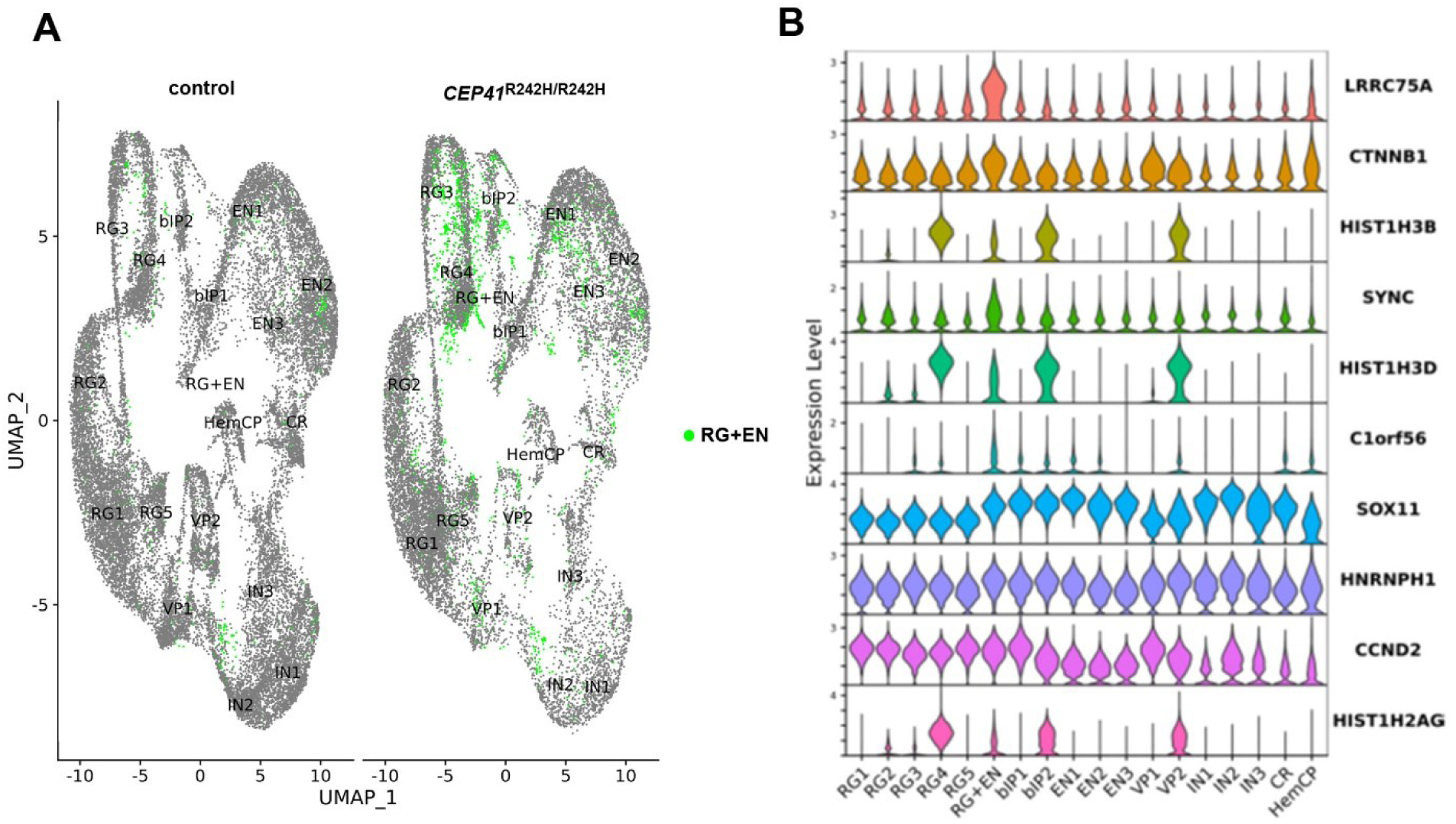
RG+EN cell type is primarily present in *CEP41* mutant organoids. (A) UMAP showing the distribution of RG+EN cells over several cell groups. (B) Violin plot revealing the expression of RG+EN characteristic markers. These cells are mainly distinguished by their high *LRRC75A* expression levels and intermediate expression of the histone markers *HIST1H3B*, *HIST1H3D* and *HIST1H2AG*.

**Supplementary Figure 5:**
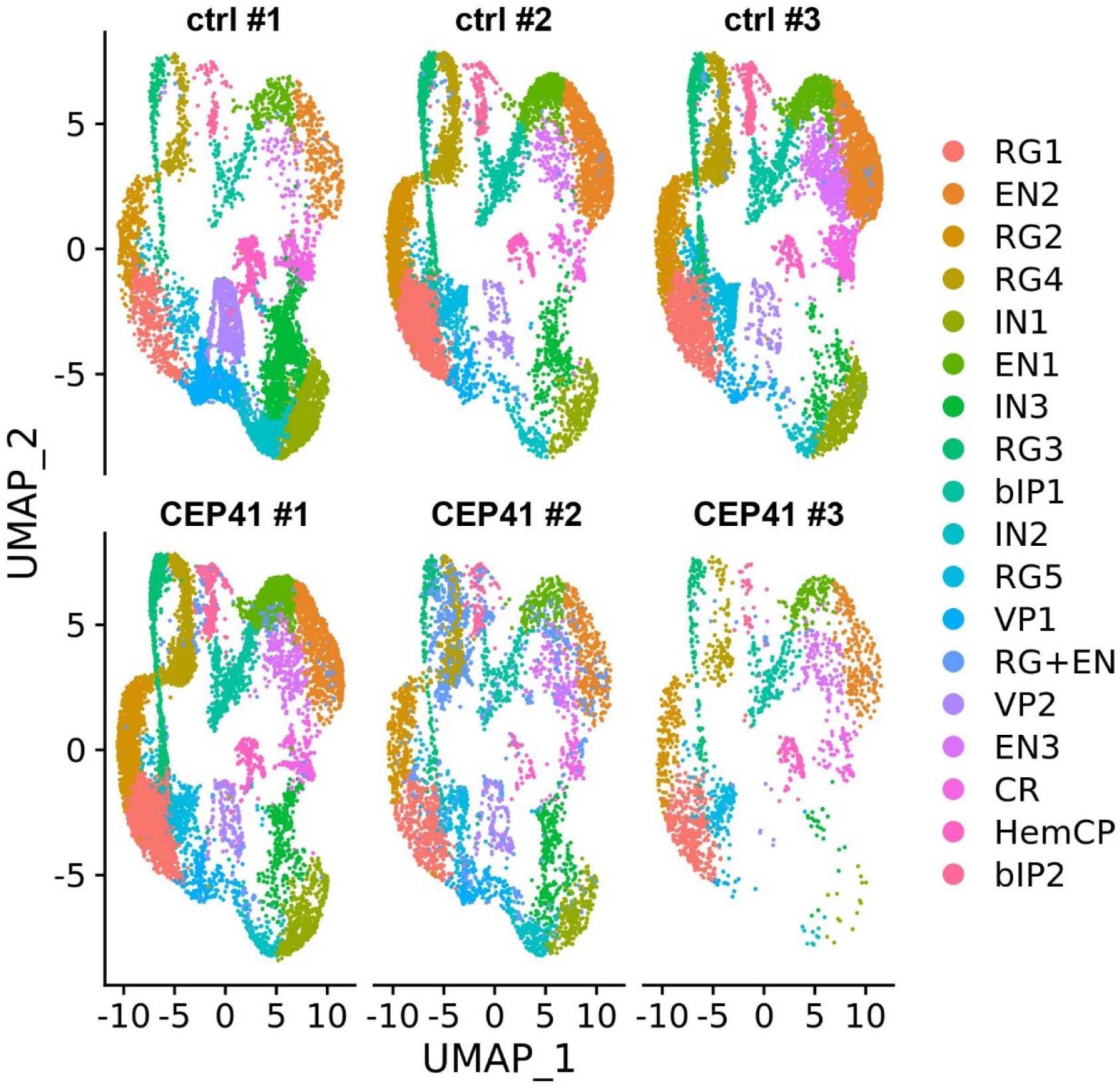
Contribution of the three control and three D38 *CEP41* mutant scRNA-seq sample to the various cell clusters.

**Supplementary Figure 6:**
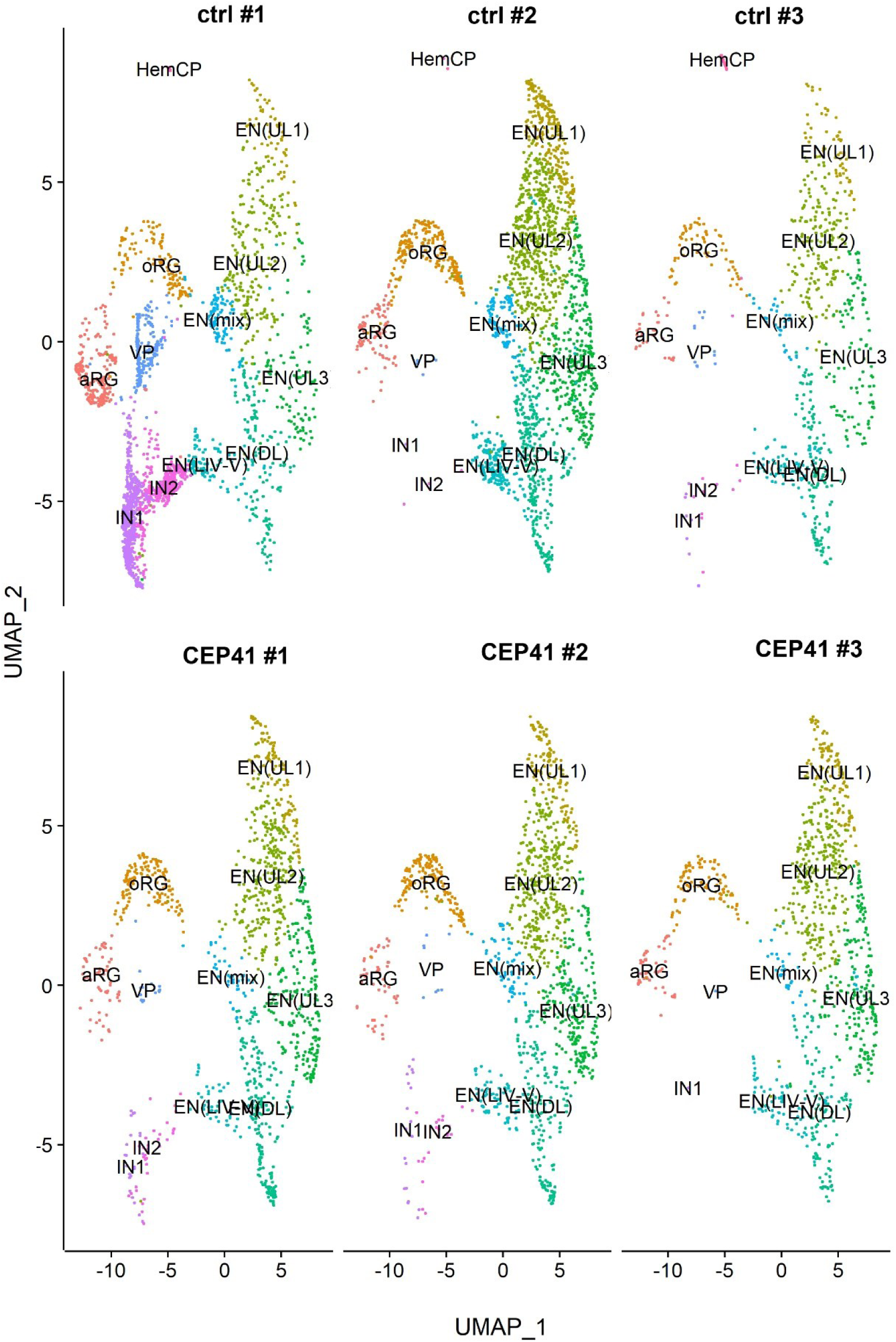
Contribution of the three control and three D94 *CEP41* mutant scRNA-seq sample to the various cell clusters.

**Supplementary Figure 7:**
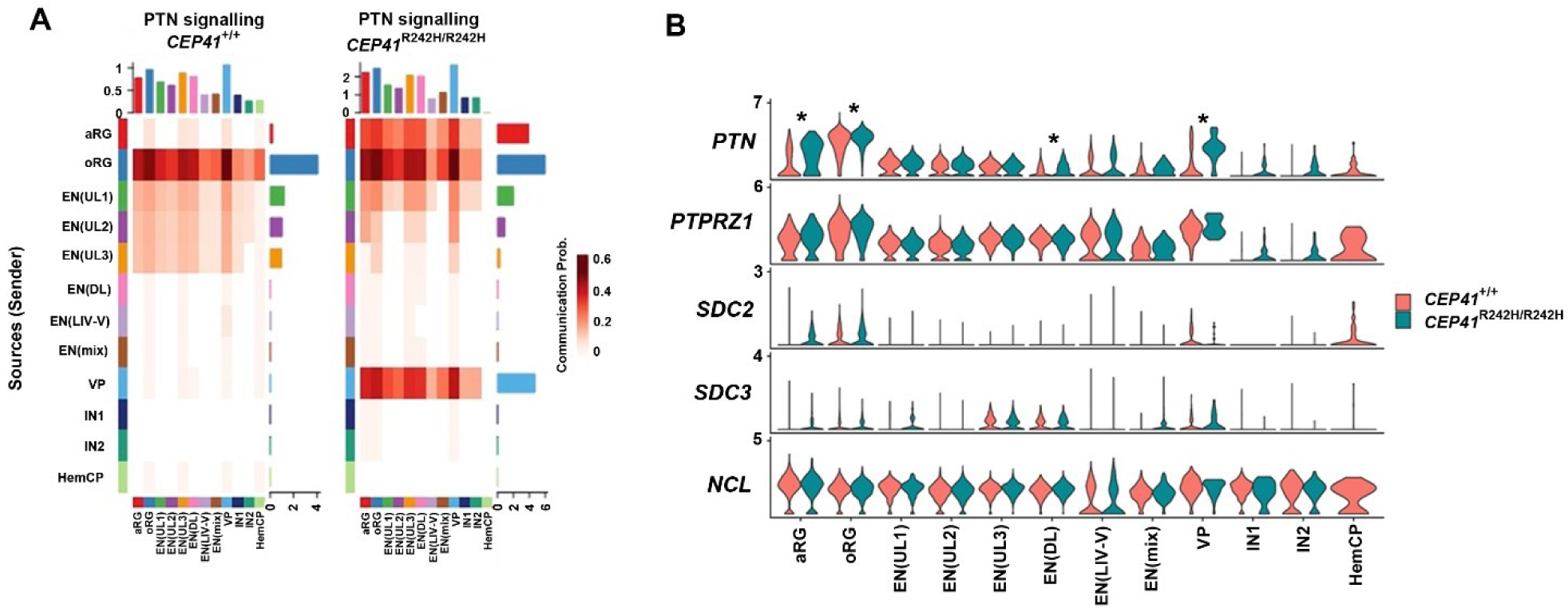
Increased PTN signalling in *CEP41*^R242H/R242H^ mutant organoids at D94. (A) Heatmap of PTN signal strength in each cell population in control and homozygous mutant organoids.(B) Violin plot showing the expression of *PTN* and its receptors in the D94 cell groups. Statistically significant *PTN* up-regulation as marked by the asterisks is observed in aRG, oRGs, EN(DL) and VP cell populations.

**Supplementary Figure 8:**
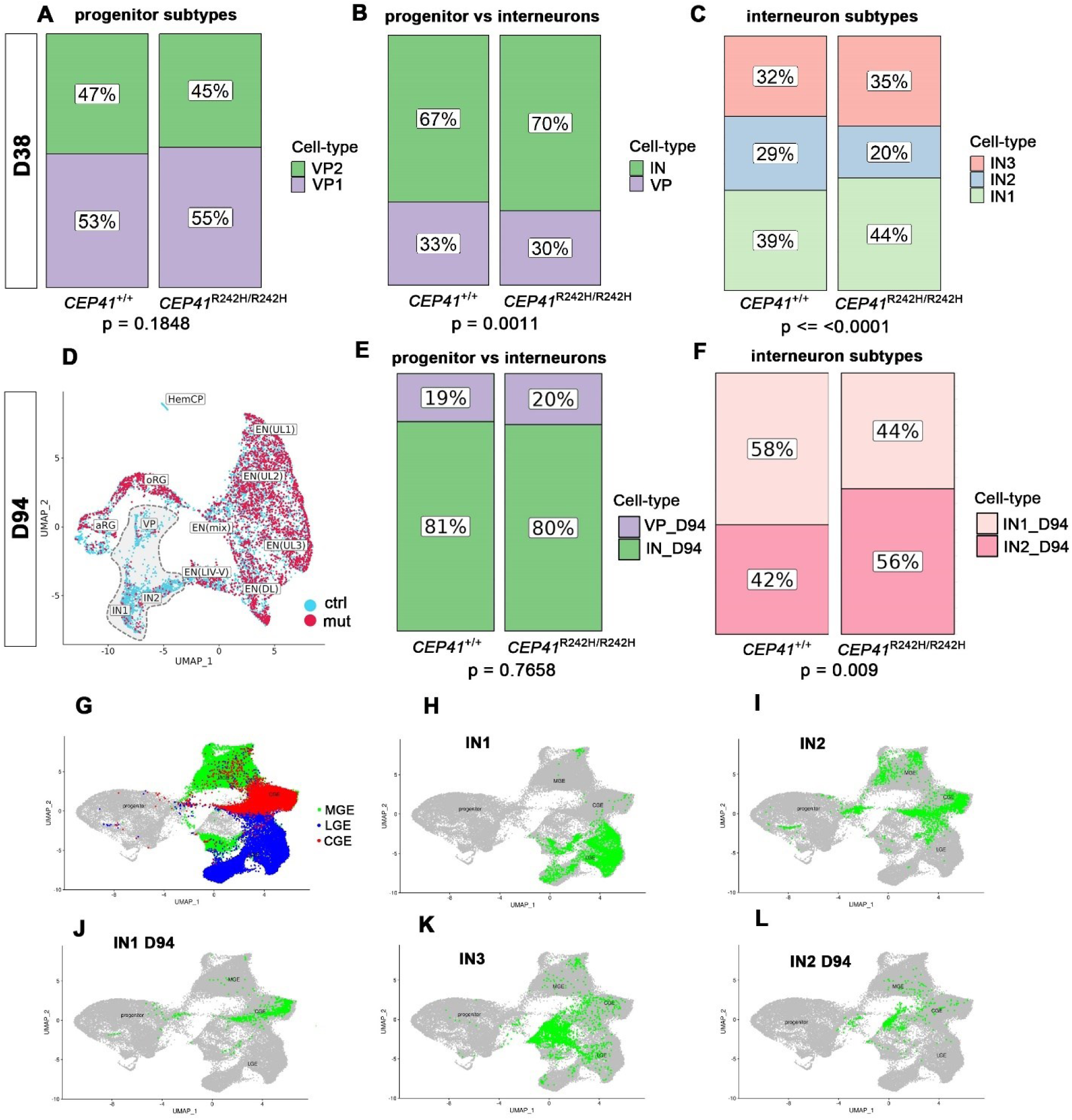
Ventral progenitor and interneuron proportions and interneuron identity in D38 and D94 organoids. (A-C) Bar plots revealing the proportions of D38 ventral progenitors and interneurons. (D) UMAP indicating the contribution of control (blue) and mutant (red) cell to each cell type at D94. Ventral cell types are highlighted in grey. (E, F) Bar plots showing unaltered proportions of D94 progenitors and interneurons (E) and a decrease in the IN1_D94 population (F). (G-L) UMAPs revealing the identity of organoid interneuron populations. Chi-square tests with (A, B, E, F) and without (C) Yates’ continuity correction.

**Supplementary Figure 9:**
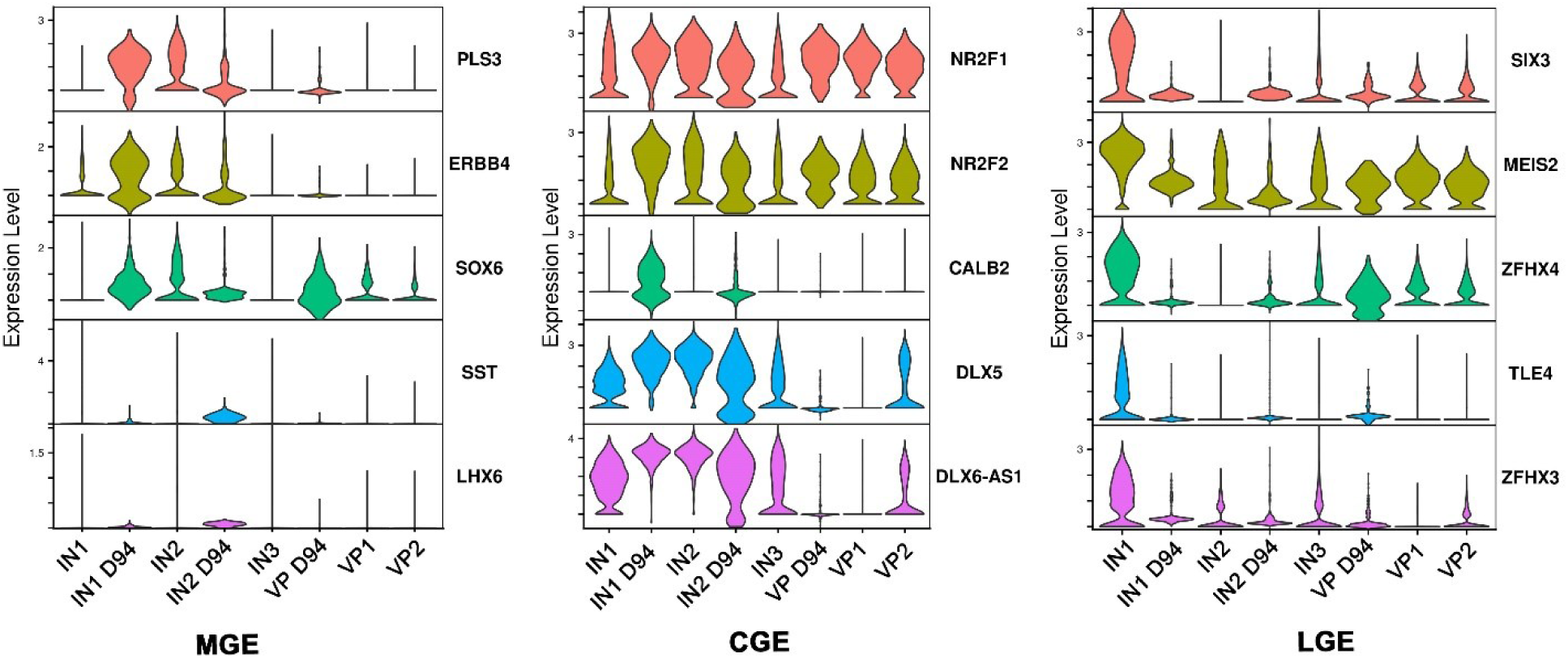
Violin plots indicating the expression levels of ganglionic eminence markers across the ventral cell types.

**Supplementary Figure 10:**
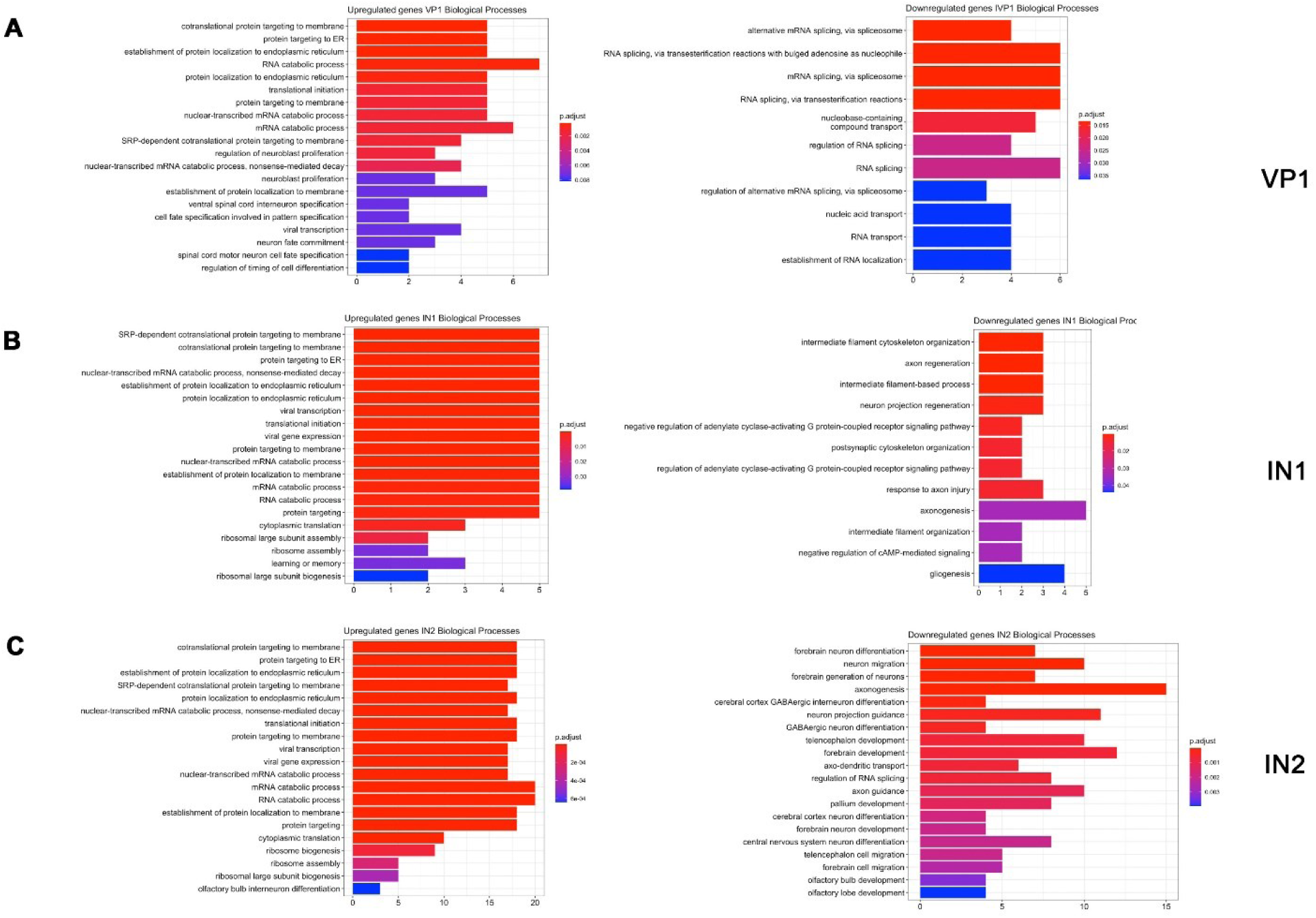
Differential gene expression in ventral cell populations of D38 organoids. (A-C) Gene ontology (GO) analysis of up-regulated (left half) and down-regulated (right half) genes in VP1 (A), IN1 (B) and IN2 (C) cells. GO terms are ordered by increasing *p*-value from top to bottom; *p*-values are shown colour coded. Count indicates the number of genes associated with each GO term.

**Supplementary Figure 11:**
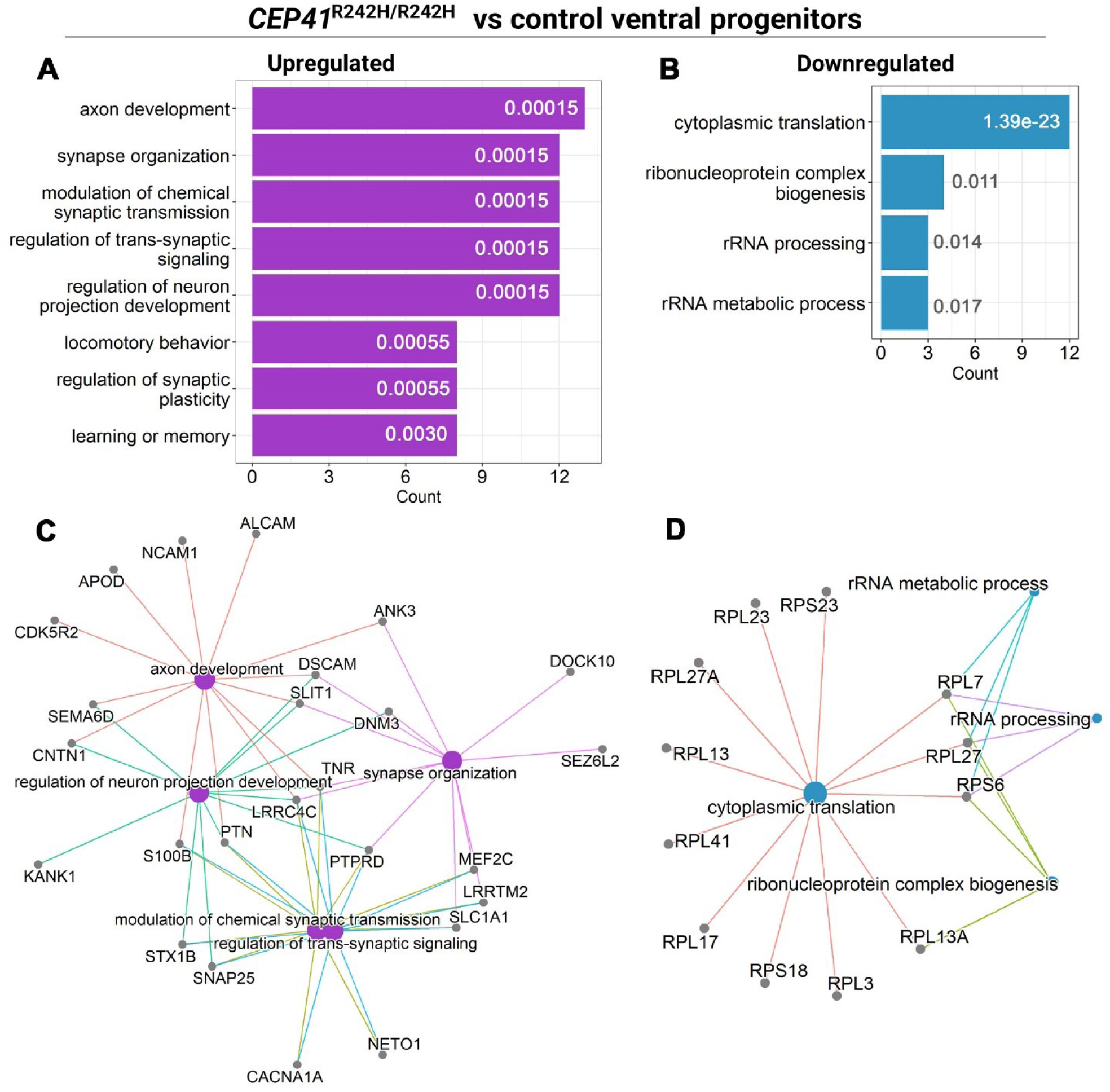
Analysis of gene expression changes in D94 ventral cell types. (A, B) Gene ontology (GO) terms enriched in up-regulated (A) and down-regulated (B) differentially expressed genes (DEGs) in D94 ventral progenitors. Least redundant out of 10 most-significant GO terms shown, ordered by increasing *p*-value from top to bottom; *p*-values are shown inside/next to bars. Count indicates the number of genes associated with each GO term. (C, D) Network analysis showing DEGs associated with the GO terms. Only significant DEGs and GO terms (adjusted *p*-value<0.05) were considered.

**Supplementary Figure 12:**
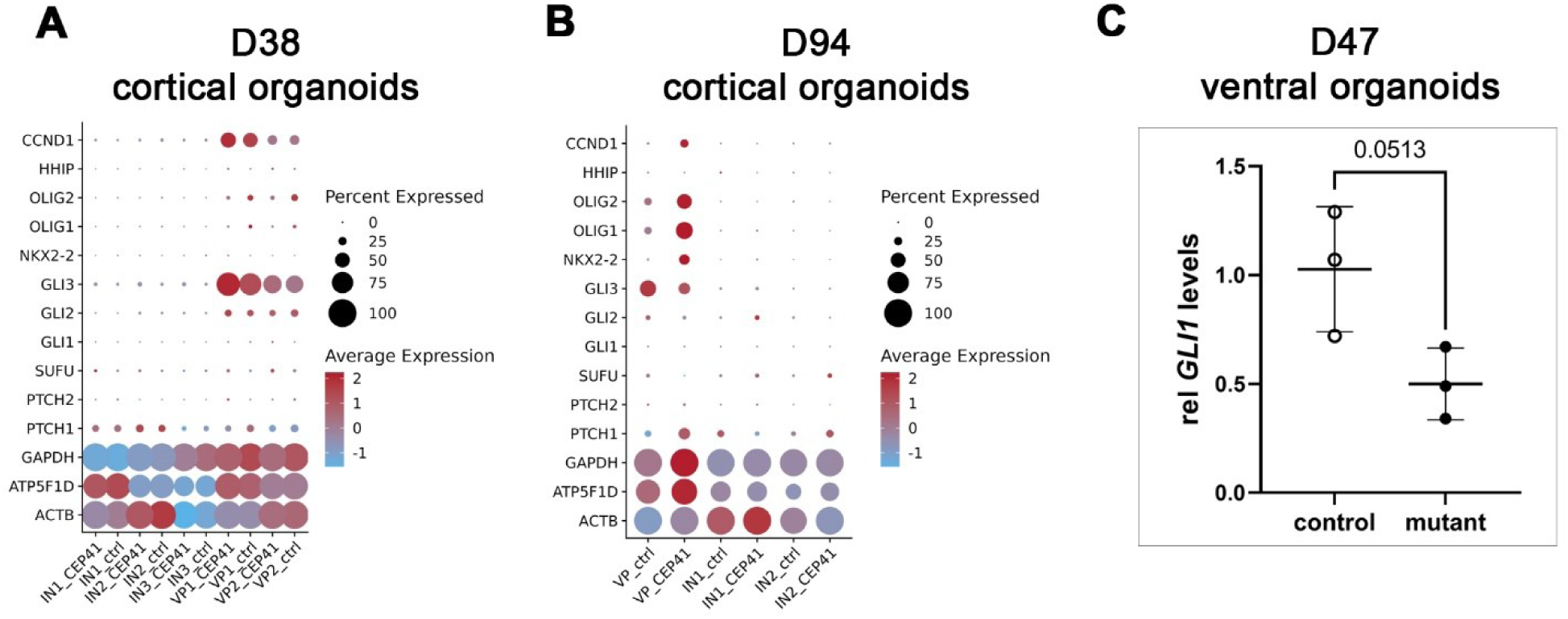
Activity of the SHH signalling pathway in *CEP41*^R242H/R242H^ mutant organoids. (A, B) Dotplots showing the average (colour) and percentage (dot size) expression of housekeeping genes and SHH signalling markers across ventral cell types in D38 (A) and D94 (B) cortical organoids. Note the low percentage expression of most analysed markers. (C) qRT-PCR analyses showing *GLI1* mRNA expression relative to *ATP5* in D47 ventral organoids. Statistical data presented as mean ± standard deviation; n=3/3 (control/mutant) lines; unpaired t tests.

**Supplementary Figure 13:**
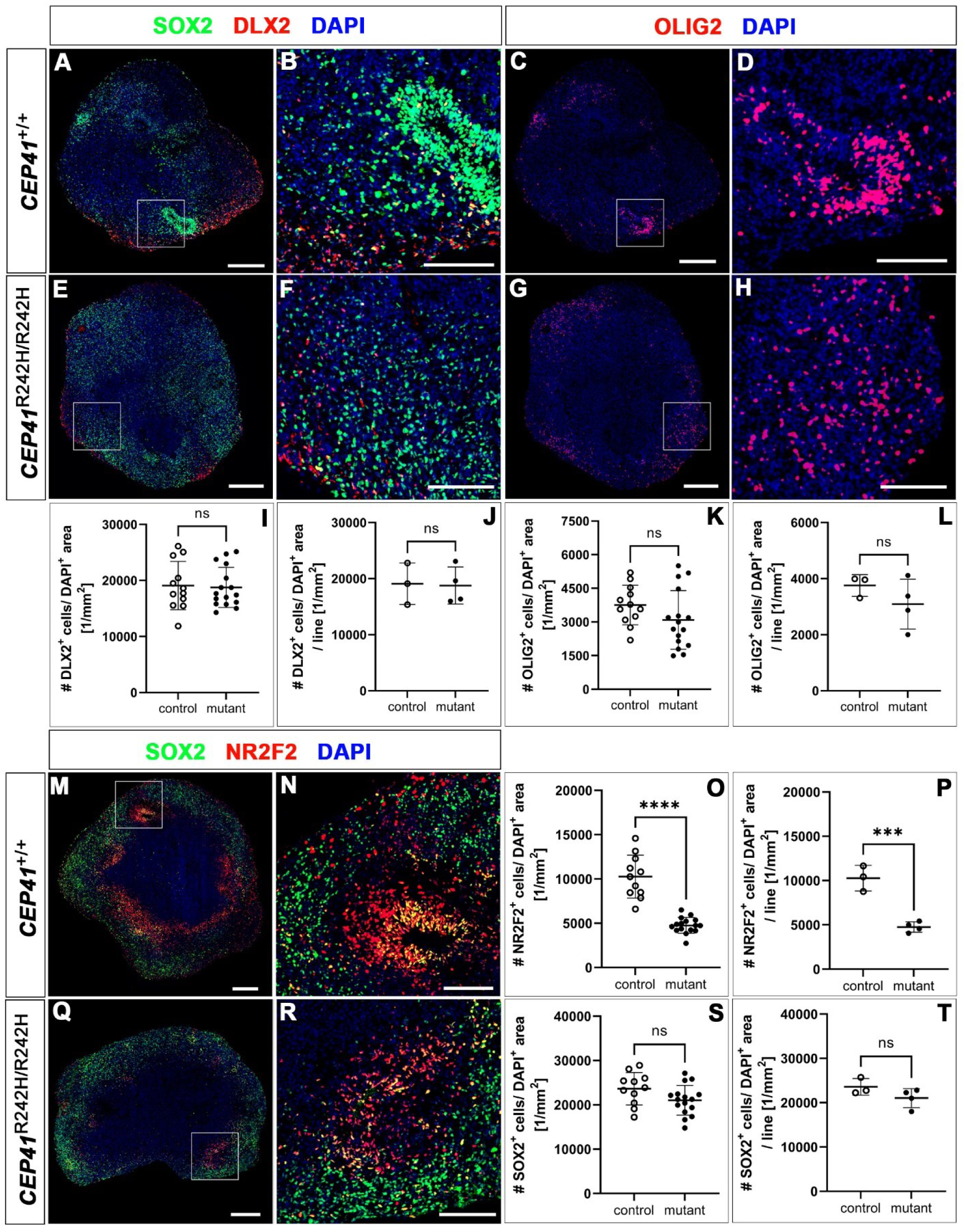
The *CEP41* mutation specifically decreases the proportion of NR2F2+ cells in ventral organoids. Immunostaining analyses using the indicated markers. **(A–H)** DLX2 and OLIG2 expression in control and mutant organoids. (I-L) Quantification of DLX2+ and OLIG2+ cell densities relative to DAPI+ area in organoids (I, J) and lines (K, L). DLX2+ and OLIG2+ cell densities showed no significant changes. (M, N, Q, R) SOX2 and NR2F2 expression. Quantification of NR2F2+ and SOX2+ cell densities in control and mutant organoids (O, S) and lines (P, T). The NR2F2+ but not the SOX2+ cell density was decreased in *CEP41*^R242H/R242H^ organoids. Statistical data presented as mean ± standard deviation (SD); control (n=11-12) and mutant (n=16) organoids; control (n=3) and mutant (n=4) lines; unpaired t tests (J-L, P, S, T); Mann-Whitney test (I, O); **** = *p*<0.0001; *** = *p*<0.001; ns = *p*>0.05.Scale bars, 250 μm (overviews) and 100 μm (magnified images).

**Supplementary Figure 14:**
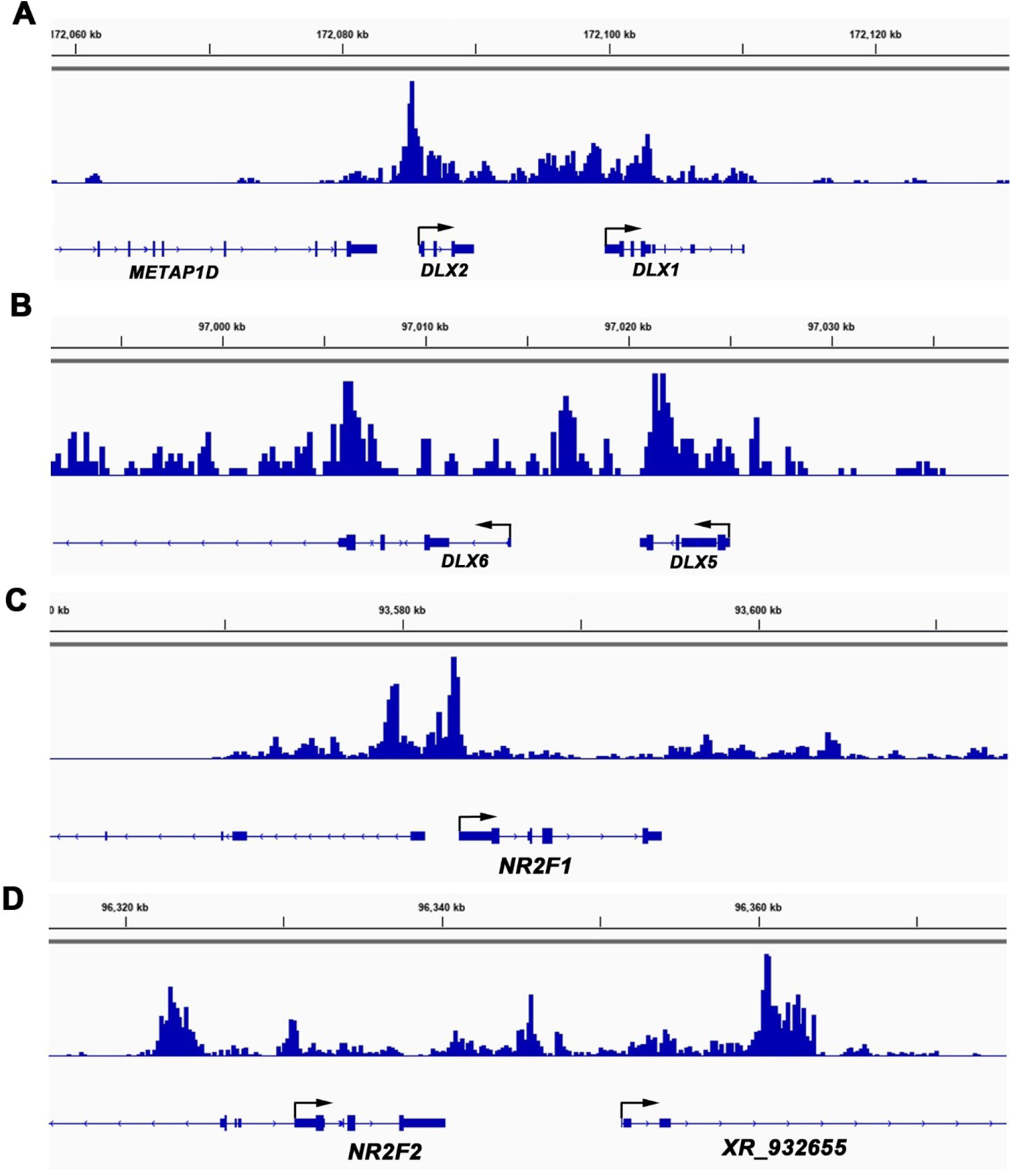
GLI3 binds to key genes controlling ventral telencephalic development. Genome browser snapshots showing GLI3 ChIP-peaks in the intergenic regions of *DLX1/2* (A) and *DLX5/6* (B), and in the promoter regions of *NR2F1* (C) and *NR2F2* (D).

**Supplementary Figure 15:**
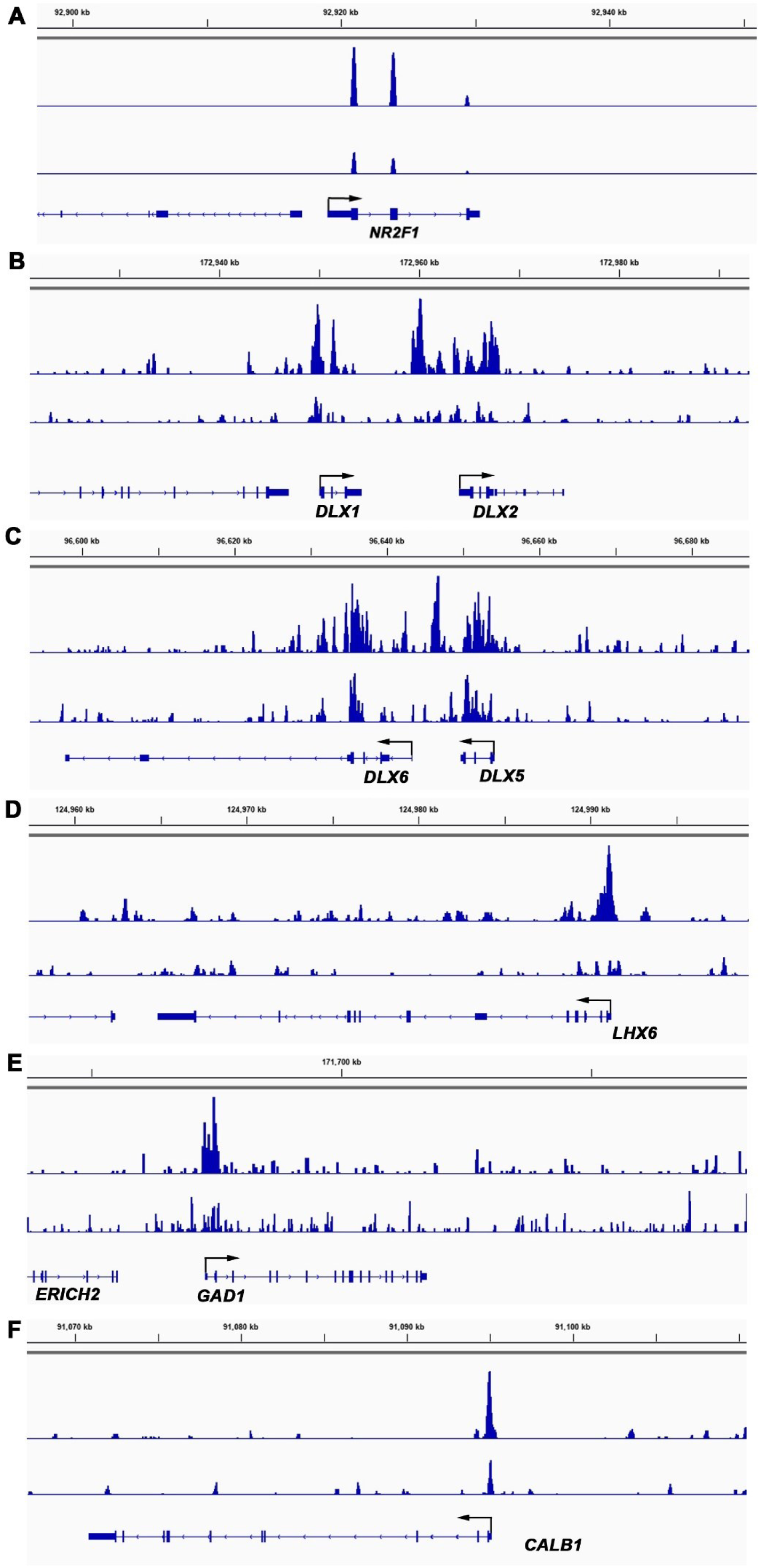
NR2F1 binds to genes controlling GABAergic interneuron development. Genome browser snapshots showing NR2F1 ChIP-peaks in the *NR2F1* promoter (A), in the intergenic regions of *DLX1/2* (B) and *DLX5/6* (C), and in the promoter regions of *LHX6* (D), *GAD1* (E) and *CALB1* (F). In each case, the upper lane shows binding of wild-type NR2F1 protein, while the lower lane represents binding of NR2F1-R112K mutant protein as a negative control.

## Notes

### Competing Interest Statement

The authors have declared no competing interest.

